# Genome-wide analysis of DtxR and HrrA regulons reveals novel targets and a high level of interconnectivity between iron and heme regulatory networks in *Corynebacterium glutamicum*

**DOI:** 10.1101/2025.01.02.631128

**Authors:** Aileen Krüger, Ulrike Weber, Julia Frunzke

## Abstract

Iron is an essential trace element required by nearly all organisms as a cofactor in enzymes, regulatory proteins, and cytochromes of the respiratory chain. Maintaining iron homeostasis is crucial, since elevated levels cause oxidative stress through the formation of reactive oxygen species. In *Corynebacterium glutamicum*, iron and heme homeostasis are tightly interconnected and controlled by the global regulators DtxR and HrrA. While DtxR senses intracellular Fe^2+^, the two-component system HrrSA is activated by heme, functioning as a global regulator of heme homeostasis. This study provides the first genome-wide analysis of DtxR and HrrA binding dynamics under varying iron and heme conditions using chromatin affinity purification sequencing (ChAP-seq). Our conditional ChAP-Seq approach revealed 25 novel DtxR targets and 210 previously unrecognized HrrA targets. Among these, *metH,* encoding homocysteine methyltransferase, and *xerC,* encoding a tyrosine recombinase, were bound by DtxR exclusively under heme conditions, underscoring condition-dependent variation in DtxR binding. Activation of *metH* by DtxR links iron metabolism to methionine synthesis, potentially relevant for the mitigation of oxidative stress. Beyond novel targets, this study highlights the interconnected nature of the DtxR and HrrA regulons, identifying 16 shared targets with in some cases overlapping operator sequences. Strikingly, we provide several examples for weak ChAP-Seq peaks, often disregarded in global approaches, that feature a significant impact of the regulator on differential gene expression. These findings emphasize the importance of genome-wide profiling under different conditions to uncover novel targets and shed light on the complexity and dynamic nature of bacterial regulatory networks.

**Importance:** The trace element iron is essential for life, but elevated levels can rapidly cause cellular damage through oxidative stress. Bacteria, like *Corynebacterium glutamicum*, tightly regulate iron and heme homeostasis via the global regulators DtxR and HrrA. This study provides the first analysis of the genome-wide binding patterns of these two regulators demonstrating significant differences in binding dependent on the tested iron regimes. Overall, we identified 25 new DtxR targets and 210 previously unknown HrrA targets, including genes with crucial roles in central metabolism and DNA repair. Notably, DtxR was shown to link iron metabolism to methionine synthesis, which might be important to protect the cell from oxidative stress. Our findings highlight the interconnected nature of DtxR and HrrA networks and underscore the value of condition-specific analysis to deepen the understanding of how bacteria adapt to environmental changes.

## 1. Introduction

Iron is a crucial trace element required by nearly all organisms due to its indispensable role as a cofactor for vital enzymes and regulatory proteins, as well as its integral presence in cytochromes involved in the respiratory chain (1, 2). Despite its biological significance, iron poses a unique challenge as it exhibits very low solubility under aerobic conditions making it a scarce nutrient in many environments. To overcome this limitation, organisms have evolved intricate measures ensuring sufficient iron supply and storage (1). However, while ferrous iron (Fe²⁺) is essential for numerous cellular processes, elevated levels can be toxic to cells. This toxicity primarily arises from its reaction with hydrogen peroxide (H_₂_O_₂_) via the Fenton reaction, which generates highly reactive oxygen species (ROS) (2). Consequently, iron homeostasis is critical for cellular fitness and is maintained by sophisticated regulatory networks integrating various parameters, such as the availability of iron, oxidative stress levels or access to alternative iron sources (e.g. heme). Notably, for the latter, the homeostasis of iron and heme is intricately interconnected in almost all organisms, which is also reflected on the level of regulatory networks.

The actinobacterial organism *Corynebacterium glutamicum* represents a model organism for studying regulatory mechanisms underlying iron homeostasis. In this species, iron homeostasis is governed by the global iron-dependent regulator DtxR and through the activity of two paralogous, heme-responsive two-component systems (TCS), HrrSA and ChrSA (3, 4). While DtxR senses intracellular Fe^2+^ levels through direct binding of ferrous ions (5), the histidine kinases HrrS and ChrS respond to heme via an intramembrane sensing mechanism (6). The DtxR protein was first discovered as “<u>d</u>iphtheria to<u>x</u>in repressor” in the related pathogenic species *Corynebacterium diphtheriae* where it controls expression of the toxin in an iron-dependent manner (7, 8). Meanwhile, DtxR-like proteins were shown to be conserved in many Gram-positive organisms and well-characterized examples include IdeR in *Mycobacterium tuberculosis* (9) or MntR in *Staphylococcus aureus* (10). In Corynebacteria, DtxR was shown to be involved in the regulation of more than 60 genes related to iron acquisition, storage as well as iron-sulfur cluster assembly (11, 12). DtxR is active as a dimer in complex with Fe^2+^ in iron sufficient conditions. Under iron limiting conditions, Fe^2+^ dissociates from DtxR rendering the protein inactive (13). Notably, DtxR also integrates into broader regulatory networks by acting as a repressor of genes encoding the key transcriptional regulators *ripA* (repressor of iron <u>p</u>roteins) and *hrrA* (<u>h</u>eme-responsive response regulator), a component of the HrrSA TCS. Besides *hrrA*, DtxR controls further genes involved in heme homeostasis, including the heme oxygenase (*hmuO*) or the heme uptake system (*hmuTUV*) (11, 14) indicating that there is considerable overlap of the iron- and heme-responsive regulons governed by DtxR and HrrA in corynebacteria.

The regulation of heme homeostasis via TCS is a widely conserved strategy in Gram-positive bacteria (15). In *C. glutamicum*, the two paralogous TCSs HrrSA and ChrSA are dedicated to sensing and responding to heme (3, 4). ChrSA is crucial for heme detoxification specifically by activating expression of *hrtBA* encoding a heme exporter (4, 16, 17). In contrast, HrrSA represents a global regulatory system orchestrating the expression of more than 200 target genes involved in heme biosynthesis, respiration and cell envelope remodeling (18).

Several previous studies focused on the elucidation of the DtxR regulon or on the control of single components providing valuable insights into its role in iron homeostasis (7, 11, 12, 19, 20). However, despite this progress, so far no study focused on the in vivo dynamics and genome-wide binding patterns of this global regulator and its potential interference with other networks. In this context, genome-wide approaches hold tremendous potential to fill these gaps by revealing the binding dynamics, structural roles, and regulatory influence of DtxR and HrrA on a global scale.

Within this study, we followed an integrative approach using <u>ch</u>romatin <u>a</u>ffinity <u>p</u>urification and <u>seq</u>uencing (ChAP-Seq) aiming to investigate binding dynamics of the two global iron- and heme-responsive regulators DtxR and HrrA, respectively. We show significant condition-specific effects on the global binding pattern leading to the identification of several novel targets, thereby connecting DtxR and HrrA to broader cellular processes including DNA topology and repair as well as oxidative stress response. Additionally, our study highlights the importance of weak – often overlooked – binding sites, which may have prominent effects on transcriptional outcomes. Consequently, our genome-wide studies provided systems level insights into the binding dynamics, interaction and interference of the two global regulators orchestrating iron- and heme homeostasis in *C. glutamicum*.

## 2. Material and Methods

### 2.1 Bacterial strains and growth conditions

The bacterial strains used within this study are listed in the supplementary Table S1. *Escherichia coli* strains for cloning purposes were cultivated in Lysogeny Broth (Difco LB, Heidelberg, Germany) media shaking at 37°C. If appropriate, 50 µg/mL kanamycin was added to the media for selection.

For a first pre-culture, the *Corynebacterium glutamicum* ATCC 13032 wild type strain or its derivatives were cultivated in liquid BHI (brain heart infusion, Difco BHI, BD, Heidelberg, Germany), inoculated from a fresh agar plate (12 mL in 100 mL baffled shaking flask for ChAP-Seq cultivation, 1 mL in deep-well plates for microtiter cultivation). Incubation followed shaking at 30°C for approximately 8 h. A second pre-culture was performed using the minimal medium CGXII supplemented with 2% (w/v) glucose (1 g/L K2HPO4, 1 g/L KH2PO4, 5 g/L urea, 42 g/L MOPS, 13.25 mg/L CaCl2 ⋅ 2 H2O, 0.25 g/L MgSO4 7 H2O, 10 mg/L FeSO4 ⋅ 7 H2O, 10 mg/L MnSO4 ⋅ H2O, 0.02 mg/L NiCl2 ⋅ 6 H2O, 0.313 mg/L CuSO4 ⋅ 5 H2O, 1 mg/L ZnSO4 ⋅ 7 H2O, 0.2 mg/L biotin, 30 mg/L 3,4-dihydroxybenzoate (PCA), 20 g/L D-glucose, pH 7.0) (21). For ChAP-seq experiments, 12 mL pre-culture 1 were transferred to 200 mL medium in a 1 L baffled shaking flask, while for the microtiter cultivation 100 µl were used in 900 µl in a deep-well plate. For the heme condition, no iron was added to this pre-culture to starve these cells from iron, while for the iron excess condition the standard amount of iron was used (36 µM FeSO4). For the iron depletion condition, a reduced amount of 1 µM FeSO4 was added to the pre-culture. Incubation was performed at 30°C shaking for approximately 16 h. From this overnight culture, the main culture was inoculated to an OD600 of 3 in CGXII with 2% glucose and either 100 µM FeSO4 (iron excess condition) or 0 µM FeSO4 but 4 µM hemin (throughout this paper further referred to as heme) (heme condition) or 0 µM FeSO4 (iron depletion condition) shaking at 30°C. For ChAP-Seq, this was cultivated in 1 L in 5 L baffled shaking flasks. For microtiter cultivation in 48-well microtiter FlowerPlates in a BioLector I microbioreactor cultivation system (Beckman Coulter GmbH, Aachen, Germany) (22) for online- monitoring. FlowerPlates were sealed with a gas-permeable sealing foil (VWR, Radnor, United States). In the microbioreactor cultivation system, backscatter (a.u.) was measured in 30 min intervals as scattered light at λ: 620 nm (signal gain: 20), while venus-fluorescence was measured at λex: 508 nm/ λem: 532 nm (signal gain: 80). Specific fluorescence (a.u.) was calculated by dividing the venus-signal by the backscatter signal for each measurement. For cultivation with plasmids, 25 µg/ml kanamycin was added.

### 2.2 Recombinant DNA work and cloning techniques

Standard molecular methods were performed according to standard protocols (23). DNA fragments were amplified via polymerase chain reactions (PCR), using chromosomal DNA of *C. glutamicum* ATCC 13032 as template and the oligonucleotides listed in supplementary Table S2. Preparation was performed as described previously (24).

Plasmids were constructed by enzymatically assembling the generated DNA fragments into a cut vector backbone using Gibson assembly (25). Sequencing of the final plasmid was performed by Eurofins Genomics (Ebersberg, Germany).

For the integration of the His-Tag and a linker (GGGS2) at the C-terminus of DtxR in the genome of *C. glutamicum*, the suicide vector pK19-*mobsacB* was used (26). Electrocompetent *C. glutamicum* cells were transformed with the isolated plasmid by electroporation (27). Then, the first and second recombination events were performed and verified as described in previous studies (28). The respective deletion was reviewed by amplification and sequencing (Eurofins Genomics, Ebersberg, Germany).

### 2.3 Reverse transcription polymerase chain reaction (RT-qPCR)

For analysis of the *C. glutamicum* strain with a tagged DtxR variant (::*dtxR*-C-linker-His), cultivation as described in 3.1 in deep-well plates was performed, harvesting cells in ice-falcons at an OD600 of 5 in exponential phase. Using the Luna One-Step RT-qPCR Kit (New England BioLabs, Frankfurt am Main) according to manufacturer’s instructions, qPCR was performed in the qTower (Analytik Jena, Jena). As a reference gene for normalization, the housekeeping gene *ddh* was used (29), besides the target gene *dtxR*. Analysis followed using qPCRsoft 3.1 (Analytik Jena, Jena) and fold-change was calculated according to the 2^-ΔΔCt^ method (30).

### 2.4 Chromatin affinity purification-sequencing (ChAP-Seq) – Sample preparation

The protocol for obtaining DNA was adapted to recent studies (18, 31). The strains *C. glutamicum* ATCC 13032::*dtxR*-C-linker-His and *C. glutamicum* ATCC 13032::*hrrA*-C-twin-Strep were cultivated as described in 3.1. After 2 h cultivation at 30°C in a rotary shaker, cells were harvested (5,000 x *g*, 4°C, 10 min). The cell pellets were washed once with CGXII without MOPS and then the cells were incubated in 20 ml CGXII without MOPS and 1% formaldehyde for 20 min at RT to cross-link the regulator protein to the DNA. The reaction was stopped via incubation with 125 mM glycine for 5 min. Then, cells were washed three times with either TNI20 (20 mM Tris-HCl, 300 mM NaCl, 20 mM Imidazol, pH 8) or buffer A (100 mM Tris-HCl, pH 8; importantly w/o EDTA, to avoid the chelation of iron ions required for DtxR activity) and the pellets were stored overnight at -80°C. For cell disruption and purification, the pellets were resuspended in approximately 20 ml TNI20 or buffer A with cOmplete protease inhibitor (Roche, Germany) and 2 mg RNase A (AppliChem, Darmstadt, Germany) and disrupted at 40,000 psi using the Multi Shot Cell Disrupter (I&L Biosystems, Germany). To shear the chromosomal DNA the samples were sonified 2 x 20 s with the Branson Sonifier 250 (Branson Ultrasonics Corporation, CT, USA) and finally supernatant was collected after ultra-centrifugation (150,000 x *g*, 4°C, 1 h).

The DNA, which was bound by the His-tagged DtxR, was purified using Ni-NTA Agarose column material (Thermo Fisher Scientific, USA) according to manufacturer’s instructions to the gravity flow protocol. Washing of the column was performed using TNI20 and the tagged protein and the bound DNA was eluted with TN buffer with rising imidazole concentrations (20 mM Tris-HCl, 300 mM NaCl, 50/100/200/400 mM Imidazol, pH 8) in each 1 mL. A Bradford assay was performed to evaluate which of the eluted fractions will be pooled for further DNA preparation. In the end, these were the last three of TNI50 elution and the first three of TNI100.

The DNA bound by the twin-Strep-tagged HrrA was purified using Strep-Tactin XT Superflow column material (IBA Lifesciences, Germany) according to manufacturer’s instructions to the gravity flow protocol. Washing of the column was performed using buffer W (100 mM Tris-HCl, 150 mM NaCl, pH 8) and the tagged protein together with the bound DNA was eluted with buffer E (100 mM Tris-HCl, 150 mM NaCl, 15 mM D-biotin, pH 8).

After the purification, 1% (w/v) SDS was added to the eluted (and pooled) fractions and incubated overnight at 65°C. Digestion of the protein was accomplished with the addition of 400 µg/ml Proteinase K (AppliChem GmbH, Germany) at 55°C for 2h. Purification of DNA followed by adding Roti-Phenol/Chloroform/Isoamylalcohol (Carl Roth GmbH, Germany) in a 1:1 ratio to the samples and consequent separation of organic phase using Phase Lock Gel (PLG) tubes (VWR International GmbH, Germany) according to manufacturer’s instructions. The aqueous phase was removed and combined with 0.1 volume of 3 M sodium acetate and twice the volume of ice-cold ethanol. The mixture was then incubated at -20°C for 2 hours. This was followed by centrifugation at 16,000 x g for 10 minutes at 4°C. The DNA was precipitated by adding ice-cold 70 %(v/v) EtOH. Following and additional centrifugation, the supernatant was carefully removed. The DNA was then dried for at least 3 h at 50°C and subsequently eluted in dH2O.

### 2.5 ChAP-Seq – Sequencing and peak analysis

The isolated DNA fragments (section 3.4) were used for library preparation and indexing with TruSeq DNA PCR-free sample preparation kit (Illumina, Chesterford, UK) according to manufacturer’s instructions, leaving out the DNA size selection steps. Libraries were quantified using KAPA library quant kit (Peqlab, Bonn, Germany) and normalized for pooling. These pooled libraries were sequenced using the MiSeq device (Illumina) (paired-end sequencing, read length: 2 x 150 bases). Data analysis as well as base calling was performed with the Illumina instrument software, yielding fastq output files. Further data analysis was based and modified according to M. Keppel et al. (18). To remove PCR amplification artifacts, sequencing data were collapsed for each sample. The processed fastq files were mapped to accession NC_003450.3 as *C. glutamicum* reference genome. This was done using Bowtie2 with the following parameters: --ignore-quals --local --very-sensitive-local --rfg 9,5 --rdg 9,5 --score-min L,40,1.2 -k 8 --no-unal --no-mixed --threads 8 -I 40 -X 800 (32, 33). The genomic coverage was convoluted with second order Gaussian kernel. The kernel was truncated at 4 sigmas and expanded to the expected peak width. The expected peak width was predicted using the following procedure: (i) All peaks higher than 3 mean coverage were detected. (ii) Points at which coverage dropped below half of the maximal peak height were detected and the distance between those was set as peak width. (iii) The estimated peak width was fixed equal to the median peak width. Convolution profiles were scanned to allow identification of the regions where first derivative changes from positive to negative. Each of these regions was determined as a potential peak with an assigned convolution score (convolution with second order Gaussian kernel centered at the peak position). Filtered peaks were normalized for inter-sample comparisons and the sum of coverages of all detected peaks was negated from the total genomic coverage. This difference was used as normalization coefficient and was divided by peak intensities.

## 3. Results

### 3.1 Genome-wide profiling of DtxR and HrrA DNA-binding under different iron-conditions

To gain an integrated view on the global iron- and heme regulatory networks controlled by DtxR and HrrA, we performed ChAP-Seq experiments revealing their genome-wide binding patterns under different iron conditions. For this purpose, tagged variants of both proteins were constructed to enable affinity purification. Fusion with a twin-Strep-tag was already successfully implemented for HrrA (18). To ensure that tagging does not interfere with DtxR’s function it was necessary to include a flexible linker sequence between DtxR and the His-tag (34) (Figure S1A). Growth analysis of the strain expressing the C-terminal His-tagged DtxR variant including the linker showed that there was no growth disturbance neither under standard conditions, nor under iron excess conditions, while an N-tagged variant featured a significant growth defect (Figure S1B). Notably, also qPCR analysis confirmed that the C-terminal His-tagged version of *dtxR* is expressed at wild-type levels in the presence of iron (2^-ΔΔCt^iron excess = 1.11 ± 0.05; 2^-ΔΔCt^standard = 1.44 ± 0.15, Figure S1C).

The genome-wide binding patterns of the two regulators DtxR and HrrA were analyzed under iron excess conditions (100 µM FeSO4) with heme as alternative iron source (4 µM heme) and under iron depleted conditions (0 µM FeSO4) (Figure 1A). The general procedure is shown schematically in Figure S2. In direct comparison, it could be confirmed that HrrA binds to a higher number of genomic targets in comparison to DtxR. Overall, a very similar binding pattern was observed for growth on iron excess and heme, respectively. This is not surprising and reflects efficient iron and heme homeostatic processes stabilizing the intracellular pool of chelatable Fe^2+^ and heme. Under iron excess conditions, sufficient heme synthesis is supported (3, 35), while in iron-starved but heme-rich conditions, heme serves as an alternative iron source via heme oxygenase HmuO (36). The iron depletion condition reflects the significantly lower activity of these regulators with less iron: While HrrA still binds to several of its high-affinity targets, binding of DtxR is almost completely abolished, ultimately leading to the strong upregulation of the iron starvation response. Pearson correlation confirmed strong consistency within triplicates of the same samples, while revealing significant differences between samples obtained from iron- or heme-grown cells. This highlights subtle but significant differences in genome-wide binding patterns observed under iron and heme conditions (Figure 1B and 1C). Notably, samples obtained from cells with a native DtxR protein (without tag) grown under iron excess condition served as negative control. Here, no unspecific binding was observed (Figure S3).

**Figure 1:**
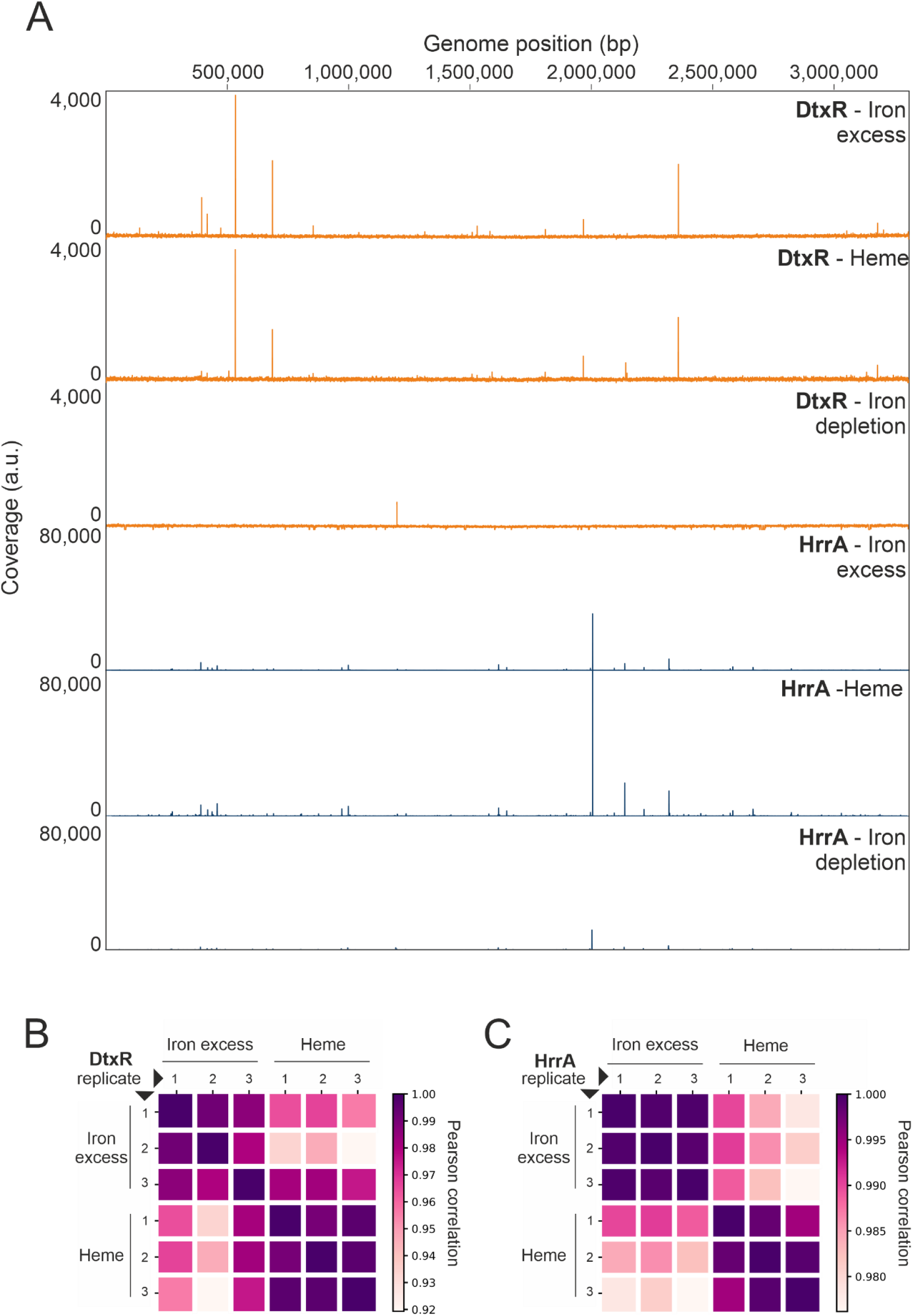
Genome-wide profiling of DtxR and HrrA DNA-binding in *C. glutamicum*. (A) Mapping of ChAP-seq reads for DtxR (orange) and HrrA (blue) to the *C. glutamicum* ATCC 13032 genome. DNA was obtained by affinity purification of DtxR and HrrA from cultures grown under iron excess (top, 100 µM FeSO4), heme (middle, 4 µM heme) and iron depletion (bottom, 0 µM FeSO4). Shown is one representative of each triplicate. Further replicates are shown in Figure S3. Note that the outstanding peak in the iron depletion condition for DtxR, found also smaller in HrrA binding, is a cryptic one and not real, resulting from technical issues as depicted in Figure S4. (B) and (C) represent the Pearson correlation of identified peaks among all replicates for DtxR and HrrA respectively binding at the two different conditions of iron excess (100 μM FeSO4) and heme (4 μM).

### 3.1.1 Condition-specific analysis revealed several new DtxR targets

For the iron-responsive regulator DtxR, overall 45 genomic targets were found across the tested conditions (Figure S5, Table S3). Representative extracted peaks for all genomic targets of DtxR are shown for the iron excess condition in Figure S6 and for the heme condition in Figure S7. Almost all are located in the upstream region of open reading frames. Among those, we could identify 20 out of the 54 targets, which were already described by previous studies using transcriptome analyses and EMSAs (11, 12). The most prominent binding peak at both iron excess and heme conditions was found upstream of NCgl0484, encoding a Fe^3+^ siderophore transport system, which was already shown be repressed by DtxR in vitro (12). Interestingly, further 25 yet unknown targets bound by DtxR were identified via ChAP-Seq (Figure S5). Those include, for example, the prophage gene NCgl1781, *metH* coding for the homocysteine methyltransferase and *cepA* coding for a putative toxin efflux permease. Among these novel targets, 11 were detected under iron excess condition and 17 targets were bound during growth on heme (Figure 2A), while only 3 targets were found in both conditions (NCgl1781, NCgl1376 and NCgl0193; genes encoding hypothetical proteins) (Table S3). Interestingly, also novel targets were found that were predicted by motif search in silico, but could not be verified in vitro in previous studies (11). These include genes encoding the putative membrane protein *wzy*, the putative oxidoreductase dehydrogenase *oxiB*, the transcriptional regulator NCgl0176 and the tyrosine recombinase *xerC*. Remarkably, we observed the trend that peaks that are closer to the transcriptional start site featured higher peak intensities (Figure 2B).

**Figure 2:**
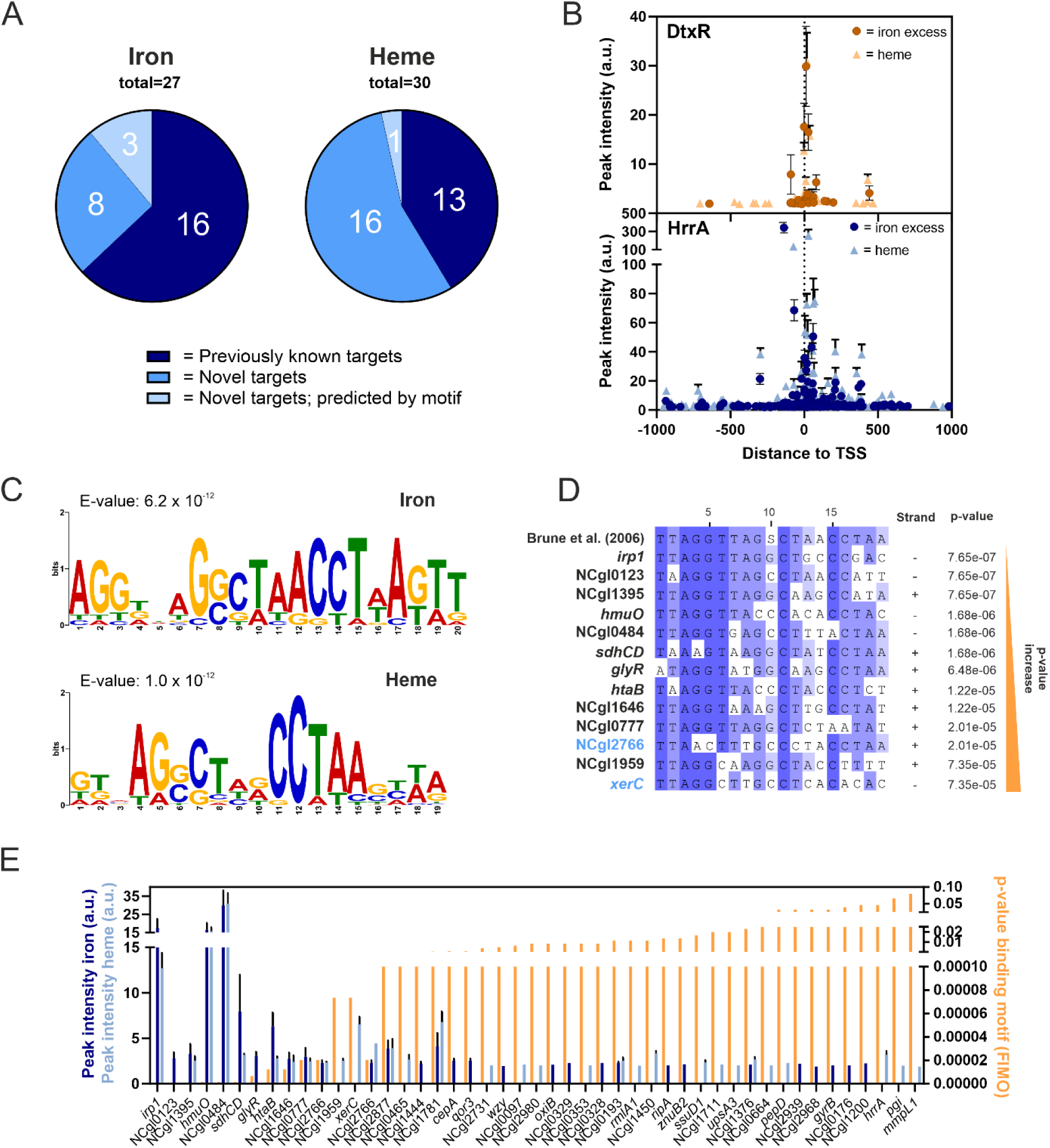
Global analysis of DtxR peaks showed correlation between peak intensity and motif conservation. (A) Pie charts comparing the number of targets bound by DtxR in this ChAP-Seq experiment that were already confirmed by previous studies (11, 12) (previously known targets), novel targets, and those that were previously predicted from motif search, but could not be found in vitro (novel targets; predicted by motif). (B) The ChAP-Seq peak intensities in arbitrary units for the iron excess (darker color) or heme (lighter color) condition were correlated to the peak distance relative to the transcriptional start site (TSS) for DtxR (blue) and HrrA (orange), respectively. (C) MEME-ChIP motif prediction of the DtxR binding motif based on ChAP-Seq binding peaks extracted from iron excess and heme conditions (37). (D) MUSCLE (Multiple Sequence Comparison by Log-Expectation) alignment (38) of previously predicted DtxR binding motif (12) and all fitting motifs found throughout the ChAP-Seq targets using FIMO (p<1.0e^-05^) (39), sorted from lowest to highest p-value. Results with a higher p-value can be found in supplementary Figure S7. Targets in light blue represent novel, so far unknown targets. (E) Correlation of peak intensities in iron excess (dark blue) or heme (light blue) condition with the p-value of the binding motif as calculated by FIMO (39).

#### 3.1.2 Condition-specific analysis of HrrA binding revealed robust homeostasis under iron excess and heme conditions

For the heme-sensitive regulator HrrA, overall 332 targets were identified (Figure S5, Table S4), with 269 binding in upstream regions and 63 inside open reading frames. 122 out of 231 targets identified by former studies could be confirmed (18). In line with previous studies, the most prominent peak for HrrA binding corresponds either to the myo-inositol-1 or 4-monophosphatase *suhB* or polyphosphate glucokinase *ppgK* (18). Additionally, 210 new targets were identified (40% in the iron excess conditions and 26% found solely during growth on heme). Among the new targets are for example the central carbon metabolism related targets *aceA, pyc* or *tkt*, as well as genes involved in signal transduction, including *cgtR2, hrcA, hspR, pdxR* or *rbsR*. In total, 83 genes are bound by HrrA only in the iron excess condition. Most targets were not identified in previous studies focusing on heme as iron source (18). This includes the genes *rpoB* encoding the beta subunit of the DNA-directed RNA polymerase, *ohr* encoding a putative organic hydroperoxide resistance and detoxification protein, the universal stress protein encoded by *uspA3* and the gene for the translation initiation factor IF-3 *infC*. Also for HrrA, we observed higher peak intensities for targets located in proximity to the TSS (Figure 2B). These results for both regulators underscore the value of conditional genome-wide analysis in providing comprehensive insights into the regulons of global transcriptional regulators.

### 3.2 Evaluation and refining of binding motifs

ChAP-Seq analysis provides genome-wide insights in binding site variations offering potential to refine motif identification. Therefore, analysis on the binding motif was accomplished using the MEME-ChIP tool (37) for both transcriptional regulators under iron and heme conditions.

Based on the previously predicted DtxR motif (TAGGTTAG(G/C)CTAACCTAA), the binding peak ChAP-Seq results for the iron excess condition lead to the motif AGGDNAGSCTAACCTWAKTT (E-value = 6.2 x 10^-12^), while the heme condition led to the similar motif GKVAGSCTHRCCTAASHWA (E-value = 1.0 x 10^-12^) (Figure 2C). The predicted motifs show a significant overlap to the previously predicted motif and fits well with the DtxR motif of *C. diphtheriae* (20). However, it became evident from our analysis that the palindromic consensus is less conserved than previously anticipated. Therefore, the predicted motif by I. Brune et al. (12) was examined within the DtxR target sequences via FIMO analysis tool (39). The alignments (Figure 2D) revealed the presence of the palindromic sequence across targets, but with considerable variation, especially for the so far unknown targets (Figure S8).

For HrrA, the found motif from iron excess CMAMCDAAAGKTKGA (E-value = 2.4 x 10^-40^) and from heme condition CAWHCRAAAGDTKKA (E-value = 9.9 x 10^-59^) also corresponds to the motif found by (18), with an even more significant E-value (Figure S9). These findings underscore the power of ChAP-Seq for characterizing binding site variability and refining motif predictions for global transcriptional regulators under different environmental conditions.

### 3.3 Validation of novel DtxR targets

Several already known DtxR targets have been confirmed in vivo by this ChAP-Seq experiment, yielding different peak variants from strong, over medium to weak binding intensity (Figure 3A). Moreover, many so far unknown potential targets were identified. To confirm binding peaks and achieve a comprehensive understanding of gene regulation, it is essential to link DNA binding with gene expression data. This allows for the assessment of whether transcription factor binding to target sites is reflected in changes at the gene expression level. On this basis, a subset of novel targets of DtxR were chosen for further analysis, varying from peaks found in both iron excess and heme conditions, or only in one of them. For each target gene, a reporter plasmid was constructed in which the coding sequence for the fluorescent protein Venus was placed under the control of the respective DtxR- targeted promoter region. All constructs were tested in *C. glutamicum* wild type and a Δ*dtxR* deletion strain at iron excess and heme conditions (Figure 3B-D, Figure S10).

**Figure 3:**
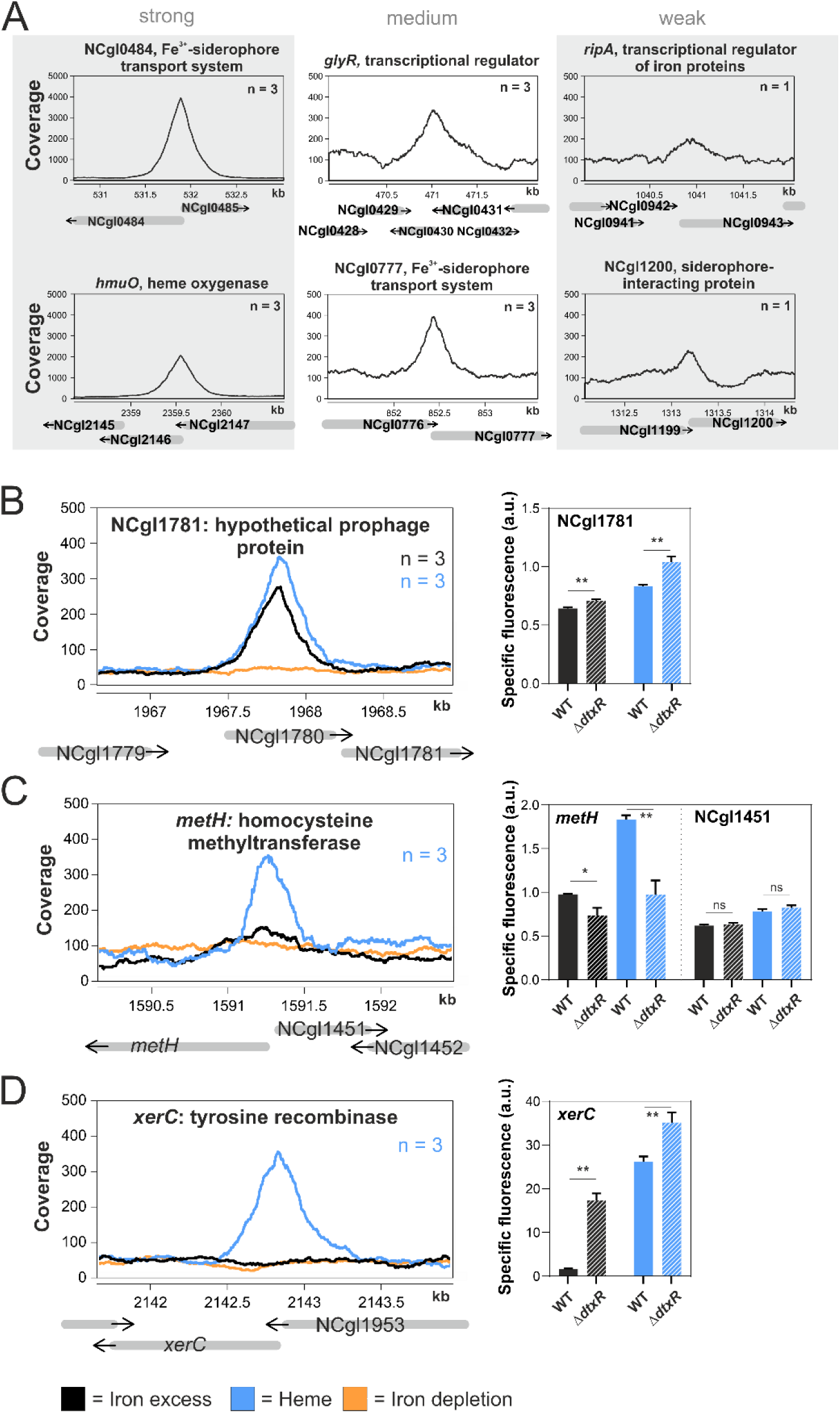
Binding peaks and reporter outputs of selected novel binding targets of DtxR. (A) Exemplarly, six known DtxR targets that were confirmed within this in vivo ChAP-Seq experiment are shown featuring strong, medium and low binding peak intensities, respectively. (B-D) Binding of DtxR to selected novel targets (NCgl01781, *metH* and *xerC*) under iron excess (black), heme (blue) and iron depletion (orange) conditions. ‘n’ represents the number of replicates where a significant peak was identified. Additionally, bar plots represent the specific fluorescent reporter output after 2h of either *C. glutamicum* WT (filled bar) or a *dtxR* deletion strain Δ*dtxR* (striped bar) transformed with a reporter plasmid coupling the activity of the respective promoter region to *venus* expression. Statistical significance was confirmed by Student’s t-test (P value ≤ 0.05).

The novel target NCgl1781 coding for a hypothetical prophage protein exhibited the highest peak intensity among novel targets and was confirmed to be repressed by DtxR under both conditions (Figure 3B). However, only minor fold-changes were observed via reporter assays. In constrast, *metH* encoding homocysteine methyltransferase was activated by DtxR in both conditions, but promoter binding was consistently observed only in the heme condition (Figure 3C). The novel target *xerC*, encoding a tyrosine recombinase, was also exclusively identified under heme conditions via ChAP-Seq but was clearly repressed by DtxR in both conditions (Figure 3D). Additional targets showed mixed responses, like *gyrB* coding for the DNA topoisomerase/gyrase IV (Figure S10A) which displayed a binding peak only in the iron condition but changes in reporter assays only in the heme condition. However, *gyrB* is likely a target underlying complex regulation involving further transcriptional regulators as well as DNA topology. In contrast, *wzy* coding for putative membrane protein was not significantly influenced by DtxR (Figure S10B). Further targets were confirmed by this approach, including *cepA* (putative toxin efflux permease), as well as NCgl0177 (putative membrane protein) and NCgl2776 (putative secreted protein) (Figure S10C-E). Overall, we identified several previously unknown DtxR targets that had not been detected under standard conditions or in in vitro binding assays.

### 3.4 DtxR and HrrA regulons are tightly interconnected

For the first time, our genome-wide profiling of HrrA and DtxR enables a direct comparison of their binding patterns on shared targets, offering new insights into potential regulatory interference. Overall, 16 targets, which were regulated by both DtxR and HrrA could be identified (Figure 4A, Table S5). Examples for binding peaks are given in Figure 4C-F with their genomic context presented additionally in Figure 4B and Figure S11. A notable example of regulatory interference between DtxR and HrrA is observed in the promoter region of the heme oxygenase *hmuO*. The locations of DtxR and HrrA binding for *hmuO* regulation are in close proximity to each other with significant overlap and their motifs separated only by 3 bp (Figure 4B-C). While DtxR acts as a repressor of *hmuO*, HrrA was shown to be crucial for activating *hmuO* expression (3). For this particular example, we observed a higher coverage of HrrA on heme and a slightly higher coverage of DtxR on iron. A similar trend – with higher coverage for DtxR on iron and HrrA on heme – was also observed for *sdhCD* (Figure 4D). Here, however, the binding sites are separated by ∼100 bp speaking against a direct interference effect. Overall, we observed a trend of higher DtxR peak intensities under iron conditions, while HrrA exhibited greater binding coverage under heme conditions (Figure S12). However, this overall trend did not establish when considering all shared targets. For example, DtxR binding to *xerC* was only observed in the presence of heme, while also HrrA showed a higher coverage under heme conditions (Figure 4E). However, this does not necessarily mean that these targets are not affected by DtxR regulation under the other condition (Figure 3D), but binding was only observable in ChAP-Seq experiments in that case.

**Figure 4:**
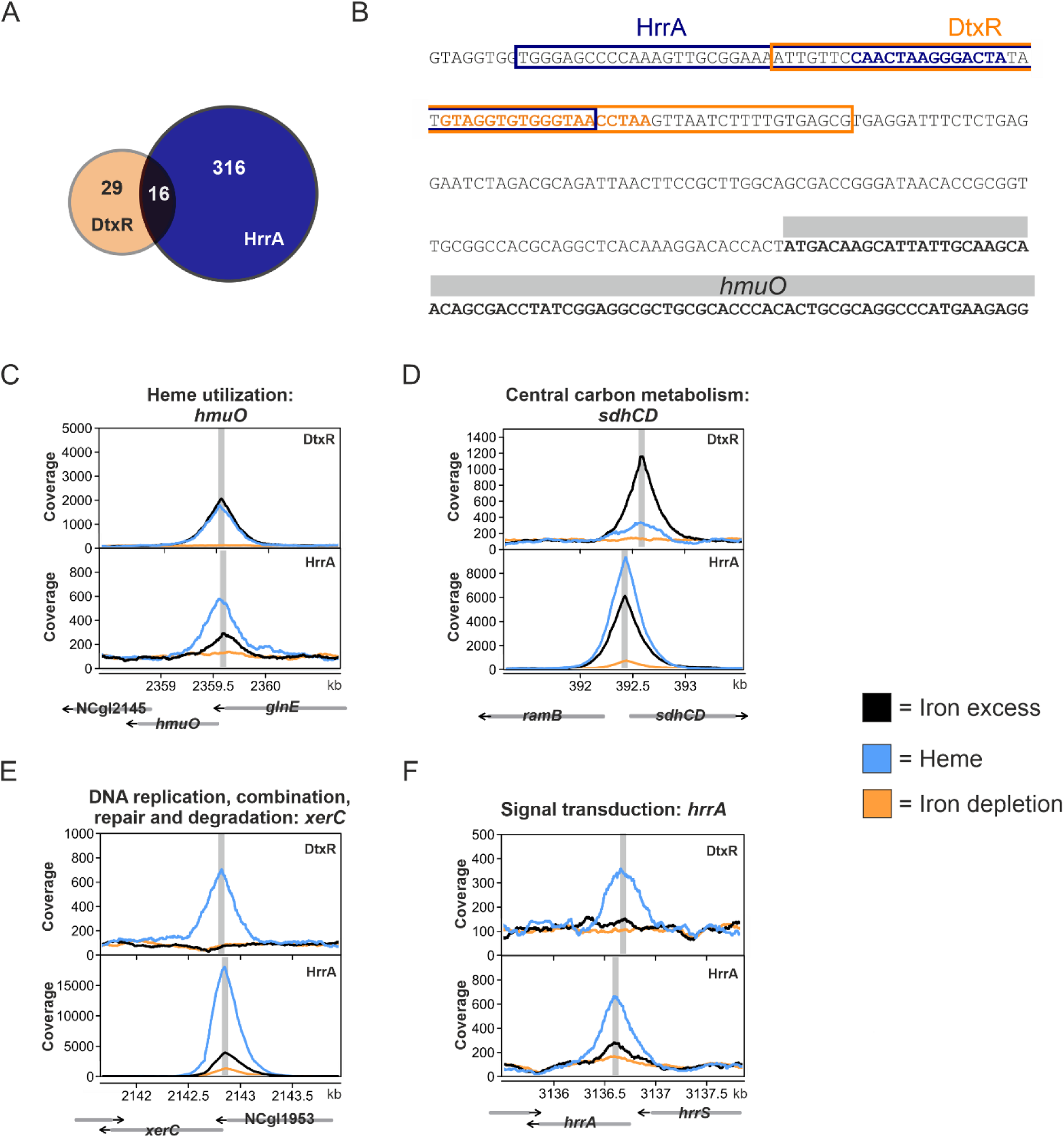
Shared targets of the regulators DtxR and HrrA show interconnection between iron and heme regulatory networks. (A) Quantitative overview of shared targets regulated by DtxR and HrrA. (B) Upstream region of *hmuO* (grey). Predicted binding regions for DtxR and HrrA are indicated by orange and blue boxes. The sequence corresponding to the actual binding motif as revealed by FIMO analysis (39) is highlighted in the respective color. (C-F) Peak detection in the region of *hmuO, sdhCD, xerC,* and *hrrA,* respectively, with gene locations represented by grey arrows. Binding peaks are shown for iron excess (black), heme (blue) and iron depletion (orange). Grey vertical lines mark the peak maxima.

Further, we could confirm binding of HrrA to itself, activating the expression of its own gene (18) (Figure 4F). While caution is necessary when directly comparing peak intensities between conditions due to differences in regulator proteins and purification methods, it was observed that HrrA binding to its own promoter increased at rising heme levels. The DtxR binding site upstream of *hrrA* is slightly upstream, but with no overlap with the HrrA motif as for *hmuO*. Surprisingly, no significant DtxR binding could be detected under iron excess, showing the high sensitivity to slight fluctuations in cultivation conditions or experimental setups.

Overall, no consistent pattern of interference between DtxR and HrrA at the promoter regions of shared targets is observed, underscoring the complexity of regulatory networks that extends beyond the interaction of just two global regulators.

### 3.5 Weak binding sites should not be ignored

ChAP-Seq experiments often produce a large number of peaks with a weak binding coverage (Figure 5A). To adress their physiological relevance, we correlated peak intensities of ChAP-Seq experiments with microarray, RT-qPCR and RNA-Seq data obtained previously (11, 12, 18). Figure 5B compares DtxR peak intensities with differential gene expression levels (Δ*dtxR*/WT) obtained from microarray analysis (11) and RT-qPCR results (12) for the targets identified in all three studies under iron excess conditions.

**Figure 5:**
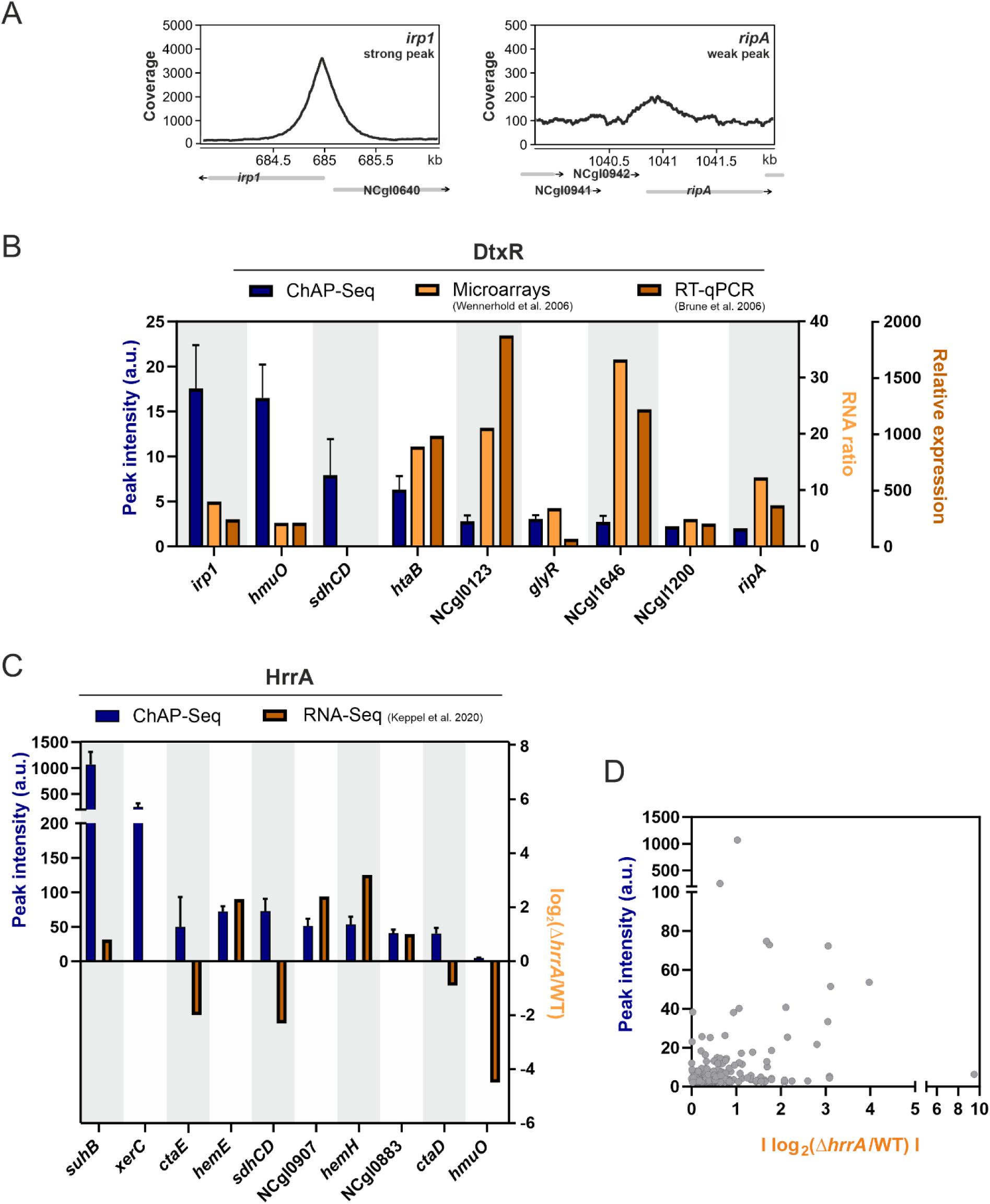
Weak binding targets should not be ignored - Correlation of ChAP-Seq peak intensity and differential gene expression. (A) Exemplarly depiction of a strong DtxR binding peak (*irp1*, iron excess condition) versus a weak binding peak (*ripA*, iron excess condition). (B) Peak intensities of the ChAP-Seq for DtxR under iron excess conditions (dark blue) were compared to the RNA ratio under iron excess (Δ*dtxR*/wild type) from a microarray analysis (11) (dark orange) as well as to the relative expression under iron excess in the deletion strain Δ*dtxR* compared to wild type analyzed via RT-qPCR (12) (light orange). (C) Peak intensities of the ChAP-Seq of HrrA in the presence of heme (dark blue) was compared to the log2(Δ*hrrA*/wild type) obtained from the time-resolved RNA-seq in the presence of heme (18) (shades of orange) for a selection of targets with highest and lowest peak intensities. (D) 2D-scatterplot showing no significant correlation between ChAP-Seq peak intensities of HrrA in the presence of heme and the absolute value of log2(Δ*hrrA*/wild type) obtained from the time-resolved RNA-seq data under heme conditions (18) for all identified targets.

Interestingly, this analysis revealed an anti-proportional trend of peak intensities and differential gene expression determined by different methods. The highest binding peaks were found in this study for *irp1* and *hmuO*, where differential gene expression levels were shown to be rather low when comparing expression in *C. glutamicum* wild type with a mutant lacking *dtxR*. Conversely, targets like NCgl0123 (hypothetical protein) and NCgl1646 (putative secreted hydrolase in the prophage CGP3 region) displayed high differential expression despite low binding coverage in ChAP-Seq experiments.

A similar trend was observed for HrrA (18), where several targets with only weak binding coverage featured strong changes at the level of gene expression when comparing *C. glutamicum* wild type with a mutant lacking *hrrA* (Figure 5C). For example, *suhB* displayed the highest binding peak but one of the lowest log2(Δ*hrrA*/WT) ratios, whereas *hmuO* showed the opposite: a low peak intensity but high differential expression in the absence of HrrA. While the correlation between peak intensities and gene expression is not consistently strong across all peaks (Figure 5D), this analysis underscores that weak binding peaks should not be dismissed as false positives to early, as they may hold significant biological relevance.

## 4. Discussion

In this study, we performed a genome-wide profiling of the global transcriptional regulators DtxR and HrrA coordinating iron and heme homeostasis in *C. glutamicum*. The obtained results provide valuable insights into the interaction and interference of iron and heme regulatory networks and demonstrate the robust homeostasis conferred by the underlying strategies. Genome-wide binding analysis revealed significant differences in binding patterns under varying iron conditions, using FeSO4 or heme as iron sources. This highlights the potential of genome-wide studies performed under different iron regimes in uncovering a substantial number of previously unknown potential targets for both regulators. The number of targets identified for DtxR and HrrA in this study aligns closely with the predicted contributions of these regulators to the *C. glutamicum* regulatory network, accounting for approximately 3% and 21%, respectively (relation percentage: 3/21 = 0.143; relation targets: 45/332 = 0.136) (40).

In this in vivo study, we successfully confirmed many previously known DtxR targets from the literature. The absence of certain targets identified in earlier in vitro studies (11, 12) may be attributed to false-negatives; however, alternative explanations are plausible. The discrepancies could arise from conditional effects, such as differences in regulatory dynamics over time, variations in cultivation conditions, or transcription factor concentrations. Recent ChIP-seq analysis of Fur, which is a functional homolog to DtxR in many Gram-negative bacteria, demonstrated a graded response of this transcriptional regulator in *Bacillus subtilis*, caused by different protein-DNA-binding affinities (41). With decreasing iron concentrations Fur derepressed its targets stepwise in ‘three waves’. Follow-up studies even determined additional targets for this well-characterized global regulator (42). This highlights the importance of binding analyses at different cultivation conditions to fully characterize the highly dynamic binding behavior of global transcriptional regulators. Another reason could be a more broadly interference with nucleoid-associated proteins or other transcription factors that mask DtxR binding (31, 43), which potentially results in observing binding to different extents at the different conditions. However, a more simple explanation could be that the other sites are too weak in binding due to e.g. hardly accessible binding sites and therefore cannot be detected with this experimental setup. Moreover, it is important to note that the HrrA analysis in this study used a different harvesting time point (2 h vs. 0, 0.5 and 4 h) (18). These differences emphasize the sensitivity of such experimental setups, particularly regarding factors like cross-linking efficiency, affinity-purification (43, 44) and condition-dependent regulatory dynamics. Several genes, including the newly identified targets *metH* and *xerC*, were identified to be bound by DtxR exclusively under heme conditions. The gene *metH* encodes the homocysteine methyltransferase, which catalyzes the last step of methionine synthesis from homocysteine. Within this study, DtxR was demonstrated for the first time as an activator of *metH* expression in corynebacteria. *metH* was already known to be regulated by the methionine and cysteine biosynthesis repressor McbR (45). For DtxR of *C. diphtheriae*, binding to methionine related genes *metA* and *mapA* was also predicted (46). Interestingly Fur, the master regulator of iron in *E. coli*, was likewise shown to regulate methionine biosynthesis via the gene *metH* (47). Methionine can function as antioxidant due to its high susceptibility to oxidation, forming methionine sulfoxide (48, 49). This oxidized form can be reduced back to methionine by methionine sulfoxide reductases (50), a process that also occurs in *C. glutamicum* via the action of the reductase MsrA (51). Subsequently, a regulatory connection between oxidative stress induced by elevated iron levels and methionine synthesis could be a valuable strategy for ROS counteraction and should be addressed in future studies. In a broader context, early studies already demonstrated that methionine promotes the catalytical activity of the heme oxygenase and ferritin in endothelial cells, thereby suppressing free radical formation (52). This connection highlights the potential for methionine to play a broader role in oxidative stress mitigation and iron homeostasis.

The repressed target *xerC*, encoding a tyrosine recombinase, was previously suggested by motif prediction, but could not yet be verified in vitro (11). Tyrosine recombinases are responsible for site-specific DNA recombination resulting in a variety of genetic rearrangements, e.g. integration, excision or inversion via breaking and rejoining single strands (53). XerC and XerD recombinases are highly conserved in many bacteria. In *E. coli,* they were shown to aid at proper plasmid and chromosome segregation during cell division, dependent on the DNA-translocase FtsK (54, 55). Contrary, the functional DtxR-homolog Fur was not described to regulate *xerC* in *E. coli* (56). The repression of *xerC* expression by DtxR suggests a potential mechanism for halting DNA recombination events under high iron conditions, possibly to prioritize DNA repair in response to iron-induced oxidative stress. Notably, *xerC* has also been shown to be repressed by HrrA, highlighting a possible convergence of regulatory control by these two global regulators (18). DtxR also controls several genes located in prophage elements (29) and XerC-like proteins were reported to play a role in phage-mediated recombination events (e.g. prophage integration). Therefore, this regulation might also link the activity of mobile genetic elements to oxidative stress levels.

Another novel target within the prophage region CGP4 is NCgl1781, which is repressed by DtxR (29, 31, 57). Previous studies identified further DtxR targets within the large prophage CGP3 (11), while it was also shown that the induction of this prophage is triggered by iron-mediated oxidative stress (29, 58). These results indicate the intricate integration of these viral elements into host regulatory networks enabling them “eavesdropping” on cellular stress responses to modulate their activity. However, many of those prophage genes regulated by DtxR encode hypothetical proteins, requiring further investigation to elucidate their specific functions and regulatory roles.

The majority of new HrrA targets has been identified under conditions of iron excess, i.e. a condition not explored in previous studies. Many of these targets are hypothetical proteins, as well as genes associated with translation and central carbon metabolism. Further validation of these targets is essential to assess the impact of this large set of targets.

Overall, we did not observe a consistent pattern of interference between DtxR and HrrA. However, the interaction between these two global iron-related regulators represents just one facet of a much more intricate regulatory network involving numerous additional factors. Recent studies in *Caulobacter crescentus* have examined the genome-wide binding pattern of IscR in comparison to Fur, the respective key regulators of iron homeostasis. These investigations identified several interference effects between these two regulators, while further suggesting the involvement of additional players (59). This is additionally highlighted by previous ChIP-Seq studies on *Mycobacterium tuberculosis*, where combination of binding profiles of in total 50 transcription factors have provided transformative insights into regulatory interferences, networks and finally pathogenesis (60). Combining such manifold genome-wide binding data with further expression analyses can significantly broaden our knowledge on regulation as demonstrated in numerous studies across domains of life (61–63).

In general, it should be noted, that binding affinities and strengths are not necessarily reflected by peak intensities (43), but further influenced by a wide range of factors in vivo, including inter- and intramolecular interactions (e.g. electrostatic or hydrophobic forces), DNA topology, and interference with other DNA binding proteins (64–66). This study further highlighted that we cannot drive simple conclusions from peak intensities to the impact of regulator binding on the level of gene expression. Weak binding peaks should not be dismissed, as they may correspond to targets that are strongly influenced by the respective transcriptional regulator.

## 5. Conclusion and future perspectives

This study provides comprehensive insights into the genome-wide binding profiles of the two global transcriptional regulators, DtxR and HrrA. Our findings highlight the potential for the discovery of novel regulatory targets through profiling under different environmental conditions, specifically using iron or heme as iron sources. The results underscore the well-orchestrated homeostasis maintained by DtxR and HrrA, with similar binding patterns observed across conditions, likely reflecting comparable but not identical intracellular pools of Fe²⁺ and heme. Importantly, our data reveal that iron-related regulatory networks are more intricate than a simple interplay between these two global regulators. The observation that targets with low ChAP-Seq peak intensities often exhibit strong regulatory outcomes suggests that even low-coverage targets should not be disregarded, as they may have significant biological relevance. In summary, the reported data expands our understanding of the global regulatory network governing iron homeostasis and emphasizes that the relation between binding behavior and functional or structural impact of prokaryotic transcription factors exists along a continuum.

## 6. Declarations

### Ethics approval and consent to participate

Not applicable

### Consent for publication

Not applicable

### Availability of data and materials

The datasets generated and/or analyzed during the current study are included in this published article and its supplementary information files. Further, all sequencing data for this study have been deposited in the European Nucleotide Archive (ENA) at EMBL-EBI under accession number PRJEB83556 (https://www.ebi.ac.uk/ena/browser/view/PRJEB83556). The custom-developed software adapted in this study for ChAP-Seq analysis is publicly available at GitHub repository under the link https://github.com/afilipch/afp.

## Competing interests

The authors declare that they have no competing interests.

## Funding

Funded by the Deutsche Forschungsgemeinschaft (SFB1535 - Project ID 458090666).

## Authors’ contributions

AK and JF wrote the manuscript. AK and UW performed experiments including cultivations, cloning, ChAP. UW performed whole genome sequencing. AK analyzed and evaluated the data. All authors read and approved the final manuscript.

## Acknowledgements

We want to thank Andrei Filipchyk for the ChAP-Seq analysis support and Max Hünnefeld for the fruitful experimental discussions.

---

**Figure S1:**
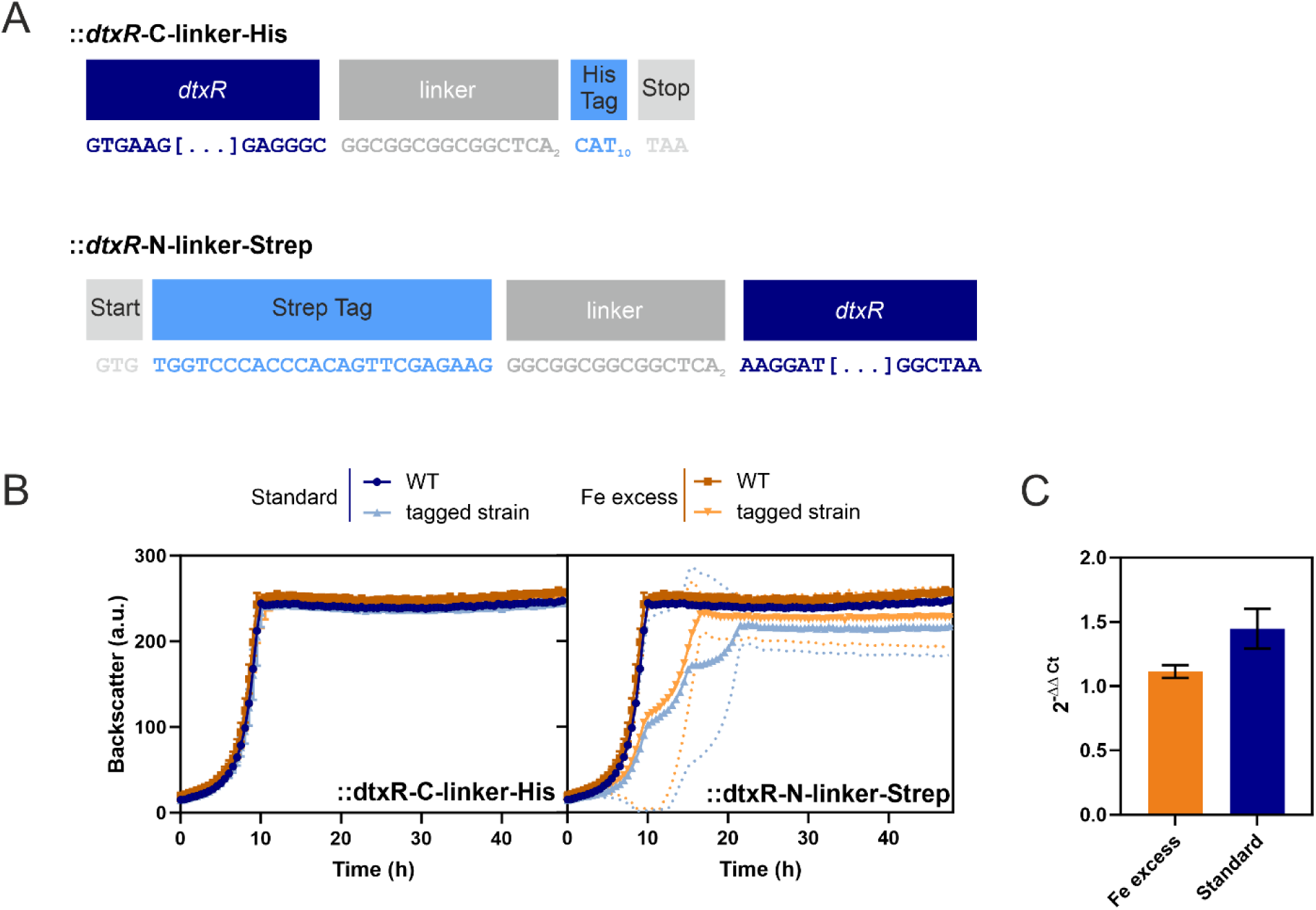
Analysis of a *C. glutamicum* strain with a tagged DtxR variant.

**Figure S2:**
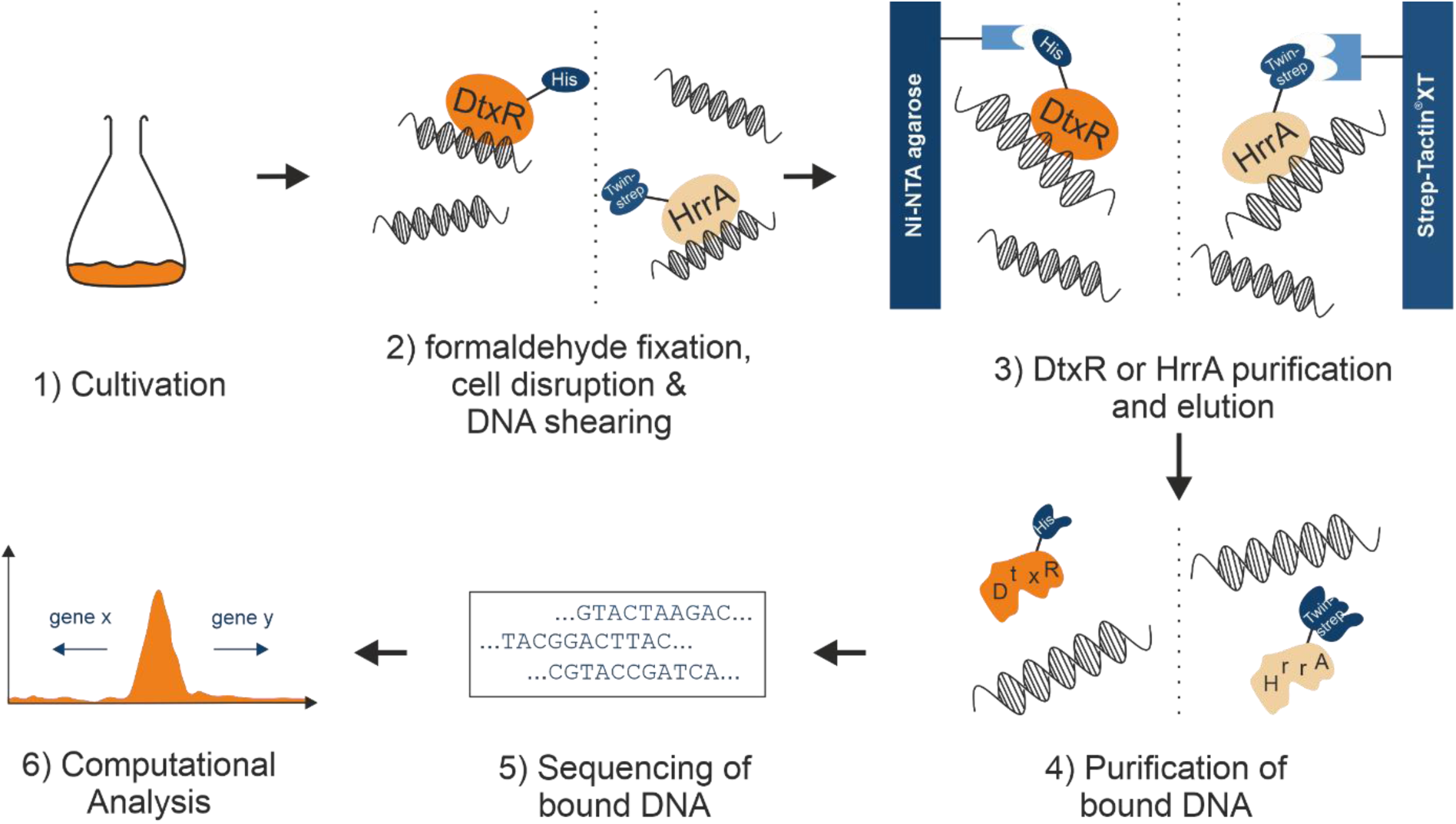
ChAP-Seq procedure for genome-wide profiling of DtxR and HrrA DNA-binding in

**Figure S3:**
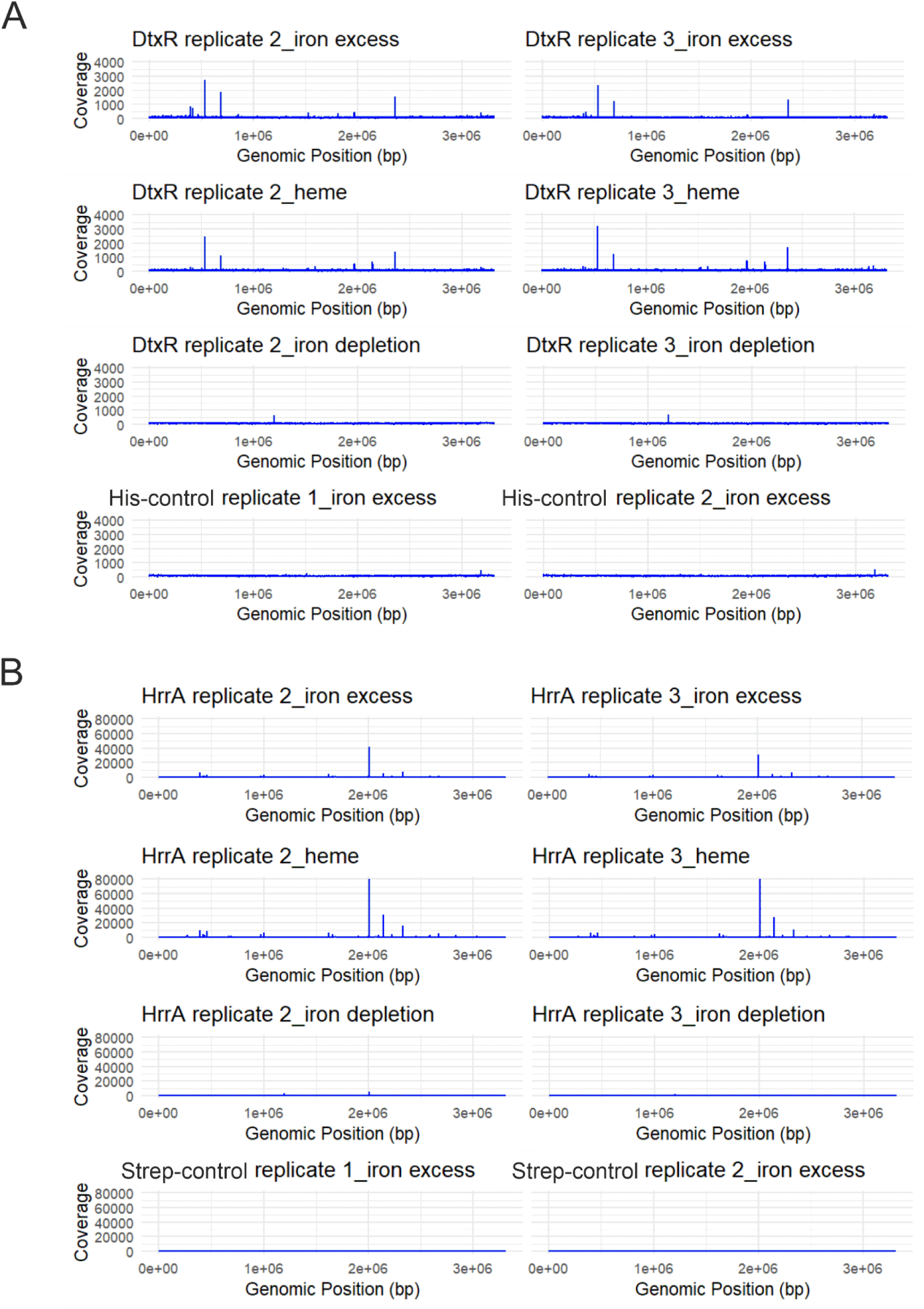
Further replicates of genome-wide profiling of DtxR and HrrA DNA-binding in

**Figure S4:**
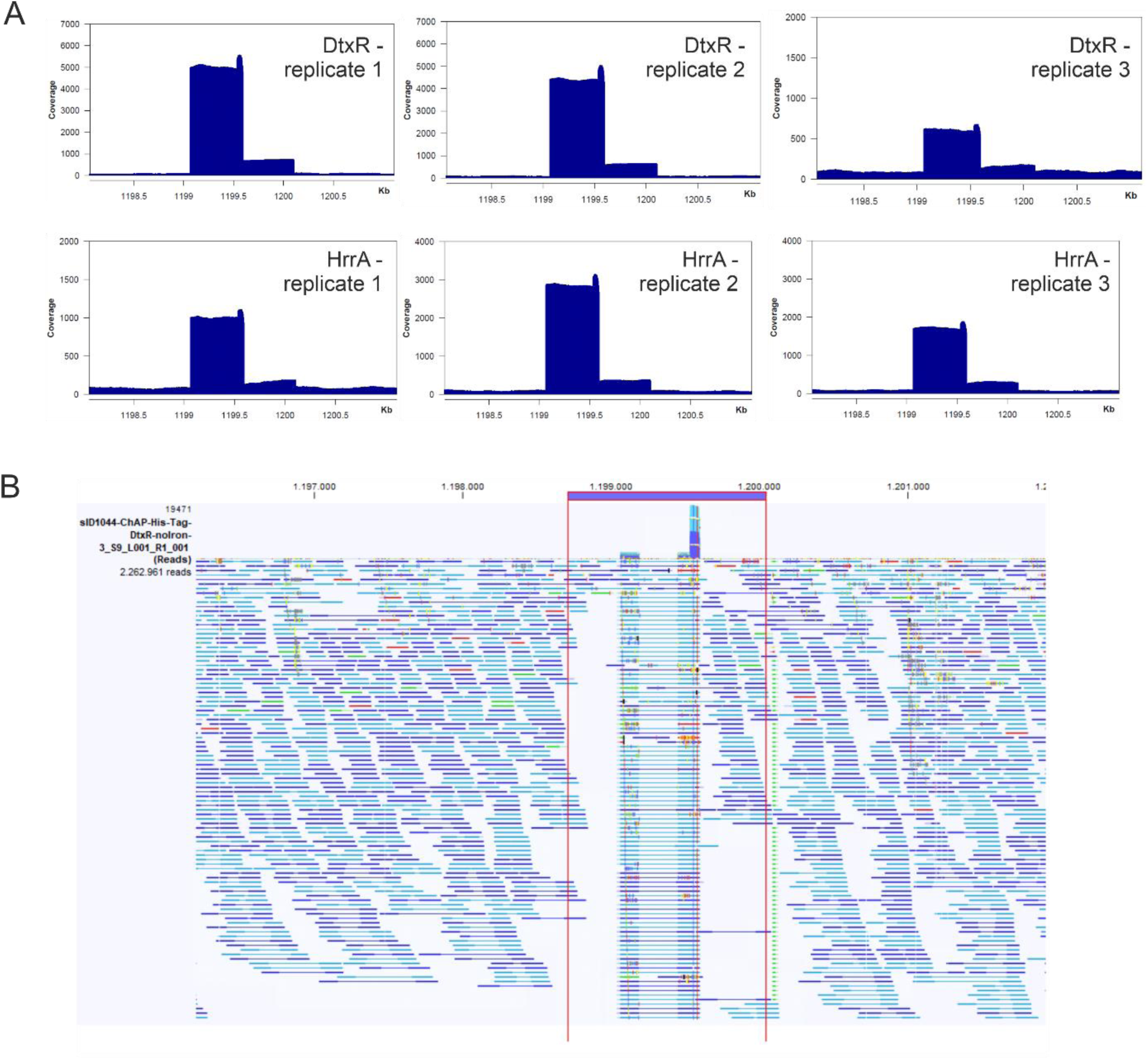
Cryptic peaks in iron depletion sequencing run.

**Figure S5:**
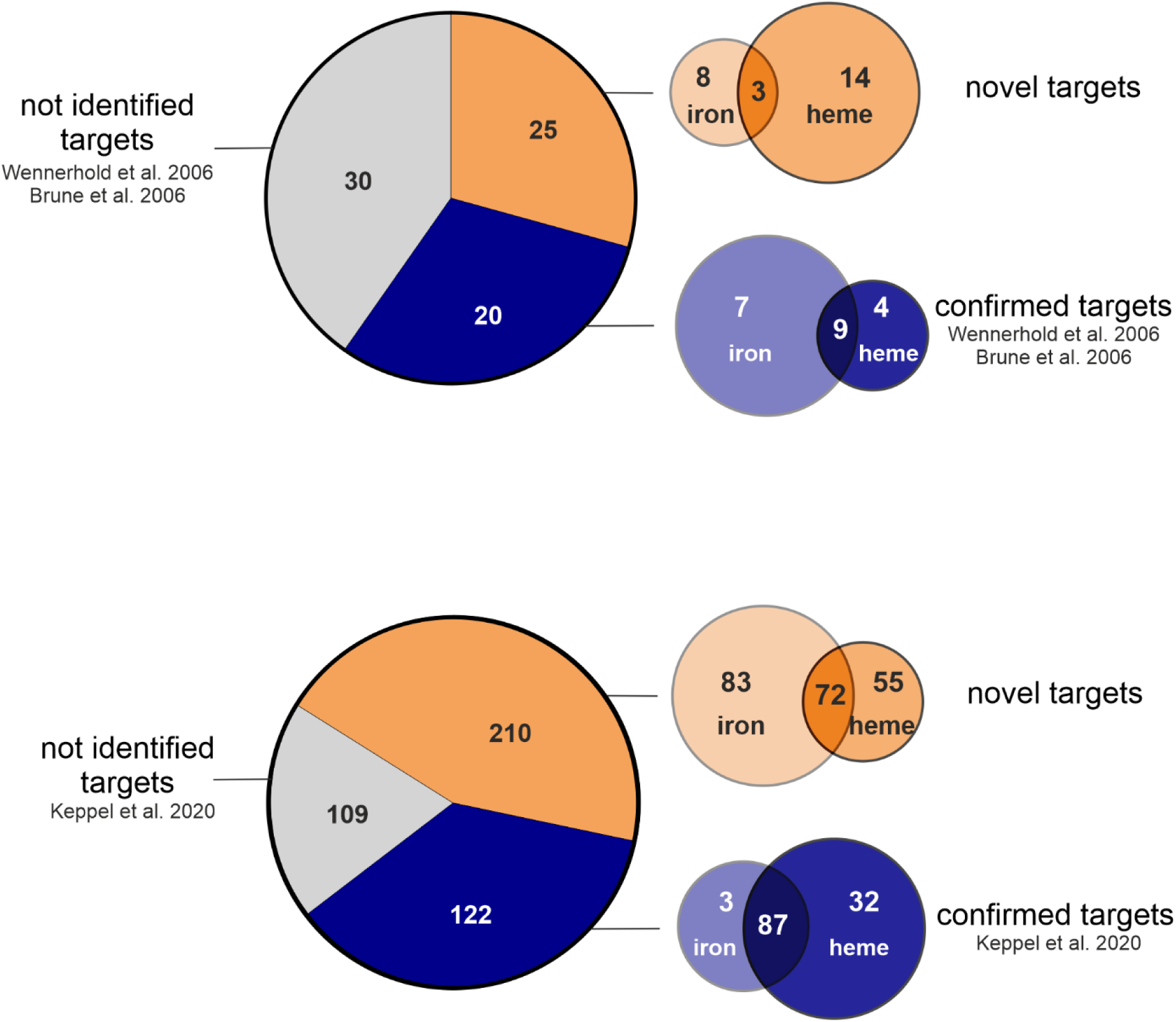
Numeric summary on targets of DtxR and HrrA.

**Figure S6:**
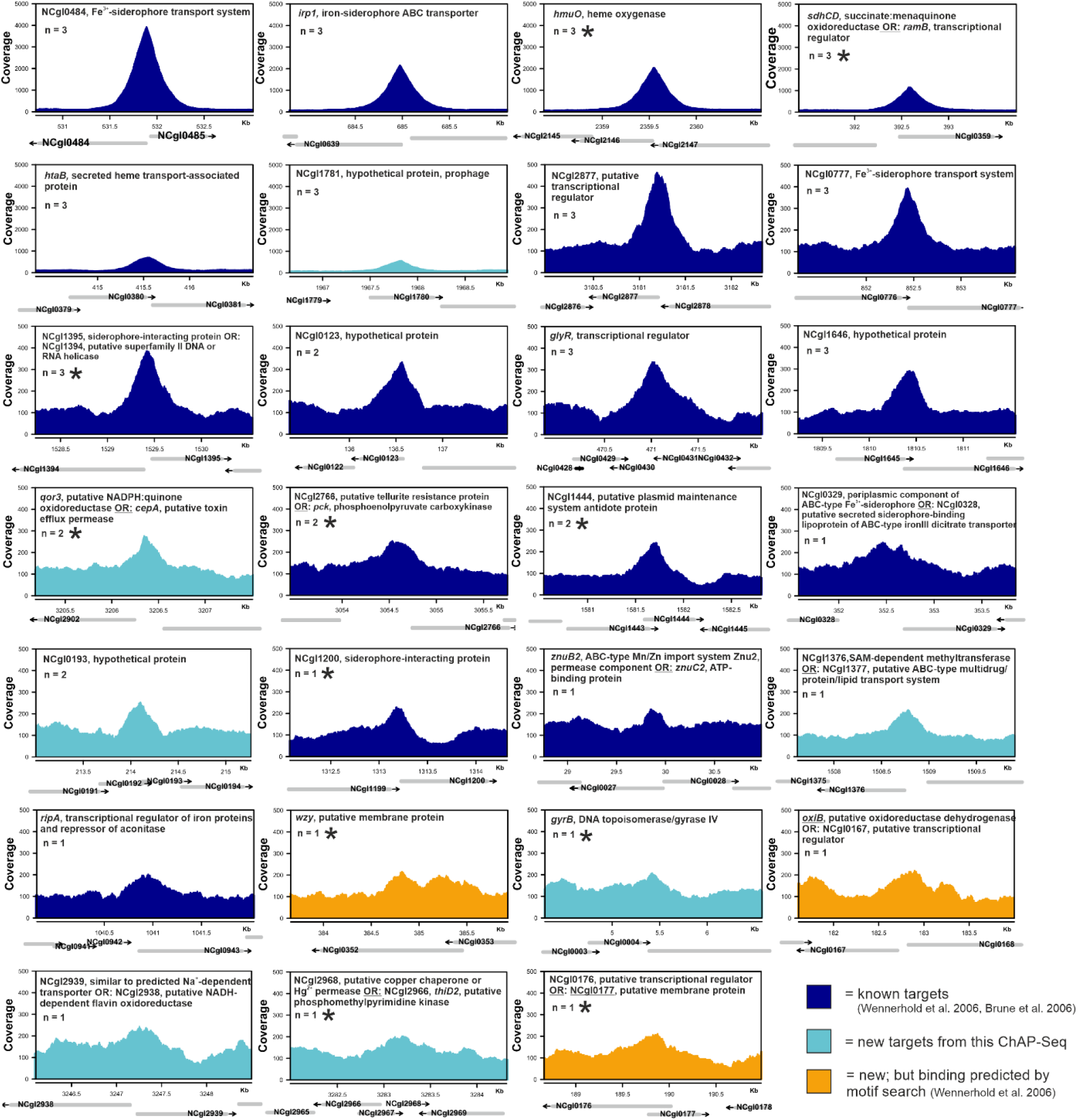
Genomic targets bound by DtxR during growth under iron excess conditions.

**Figure S7:**
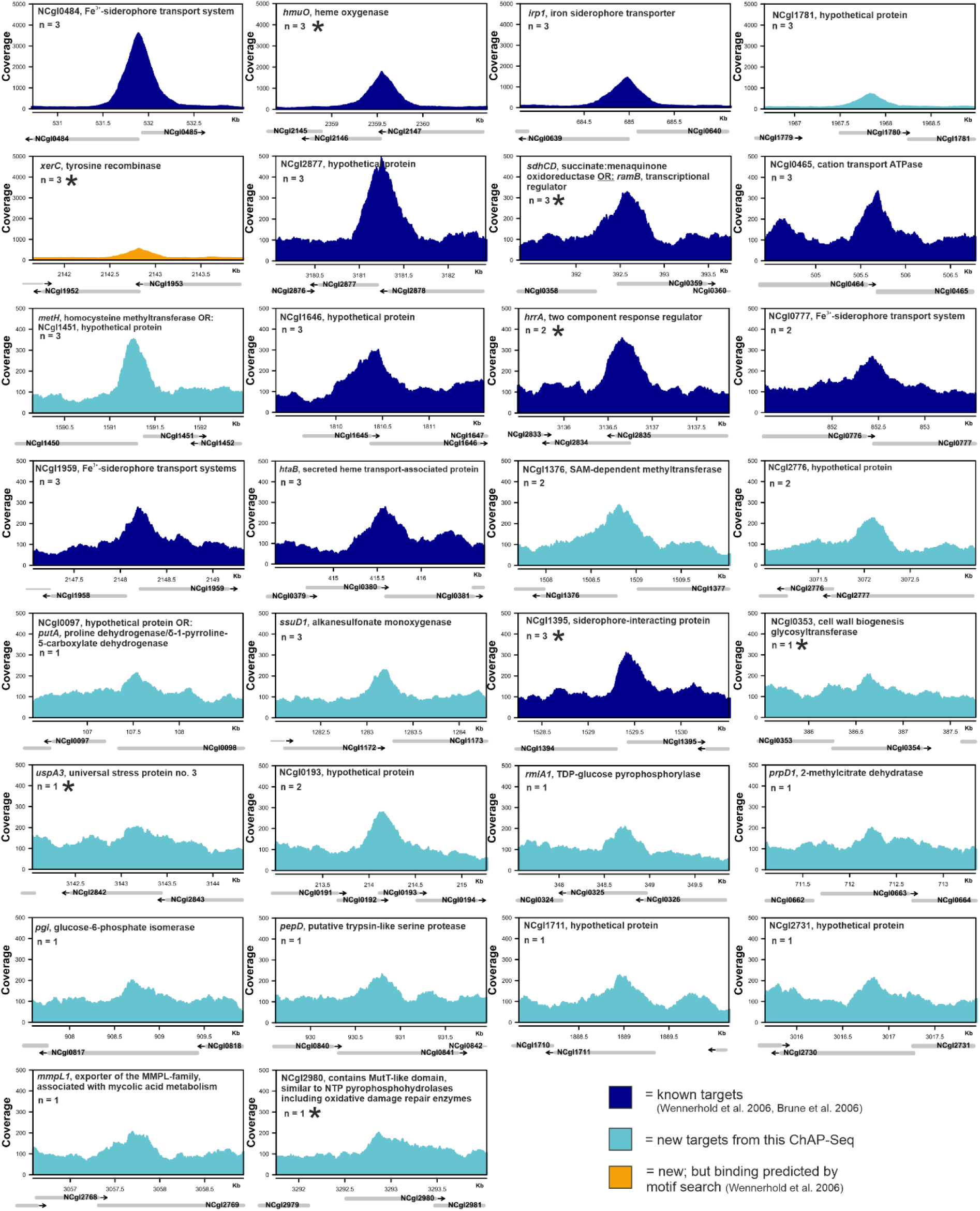
Genomic targets bound by DtxR during growth under heme conditions.

**Figure S8:**
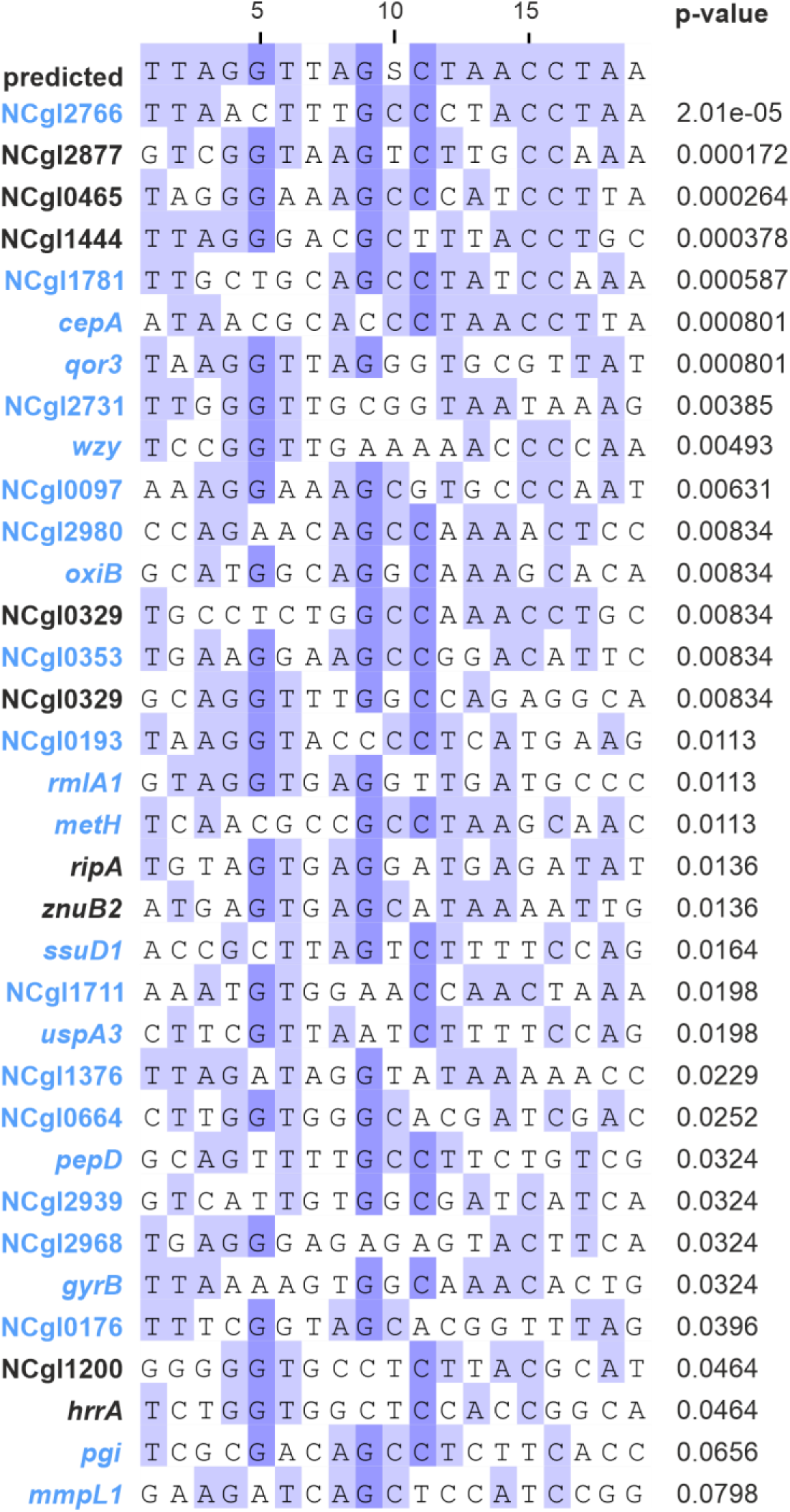
Motif alignment for DtxR binding as predicted from ChAP-Seq data (p > 1.0e^-05^).

**Figure S9:**
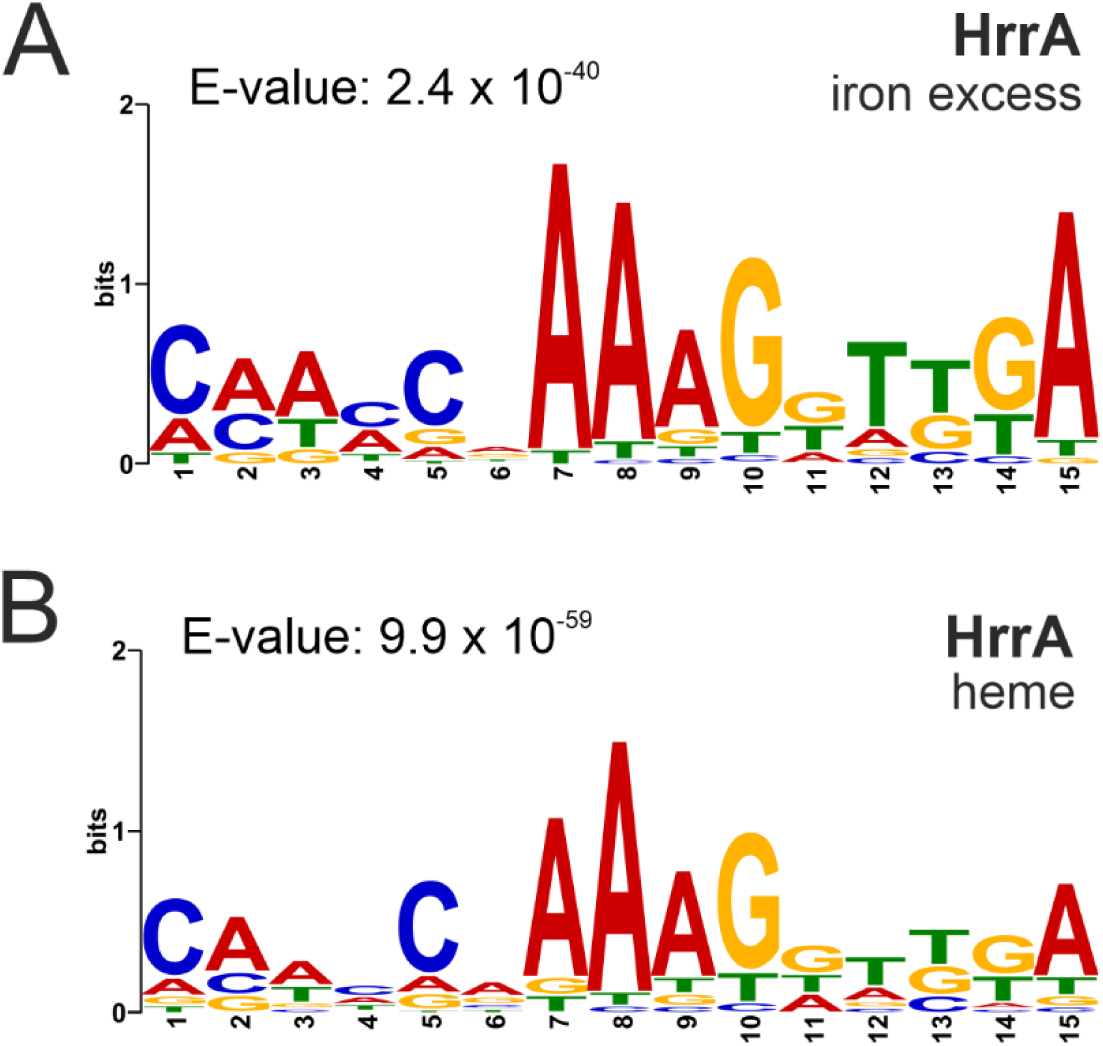
HrrA Motif predicted from ChAP-Seq results.

**Figure S10:**
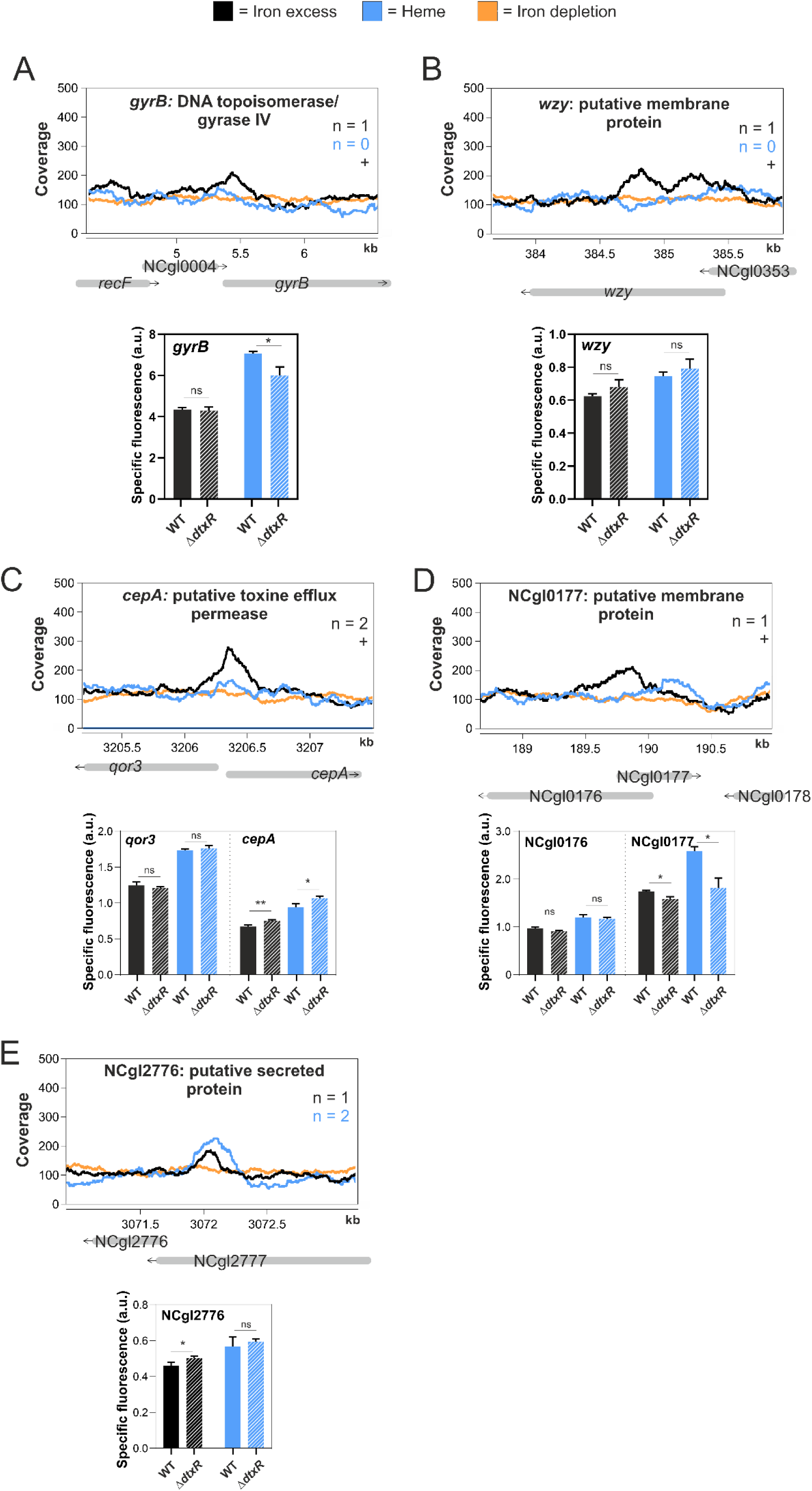
Binding peaks and reporter outputs of further selected novel DtxR targets.

**Figure S11:**
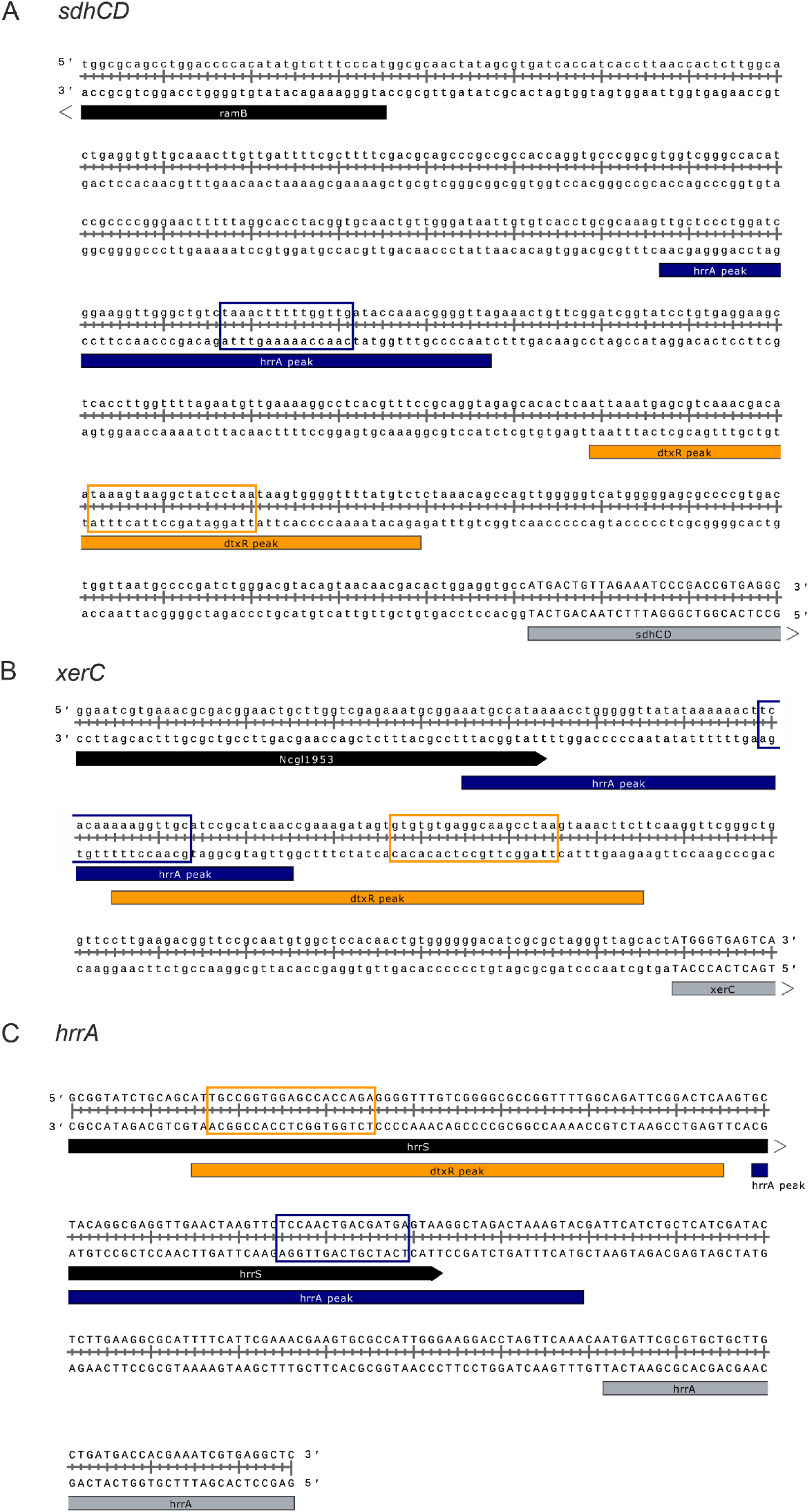
Location of DtxR and HrrA peaks in the promoter region of selected shared target genes

**Figure S12:**
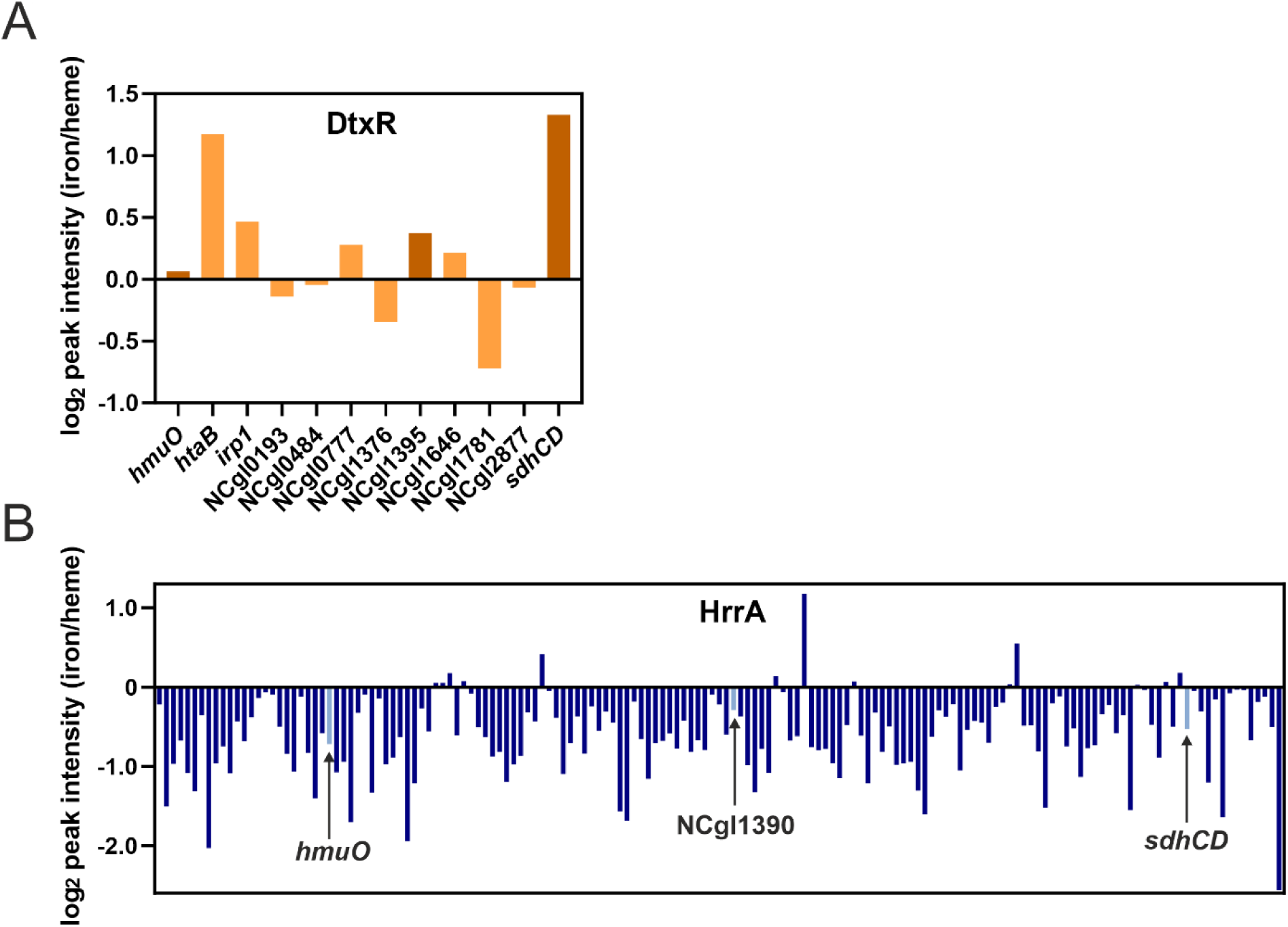
Ratios of iron and heme peak intensities.

**Table S1:**
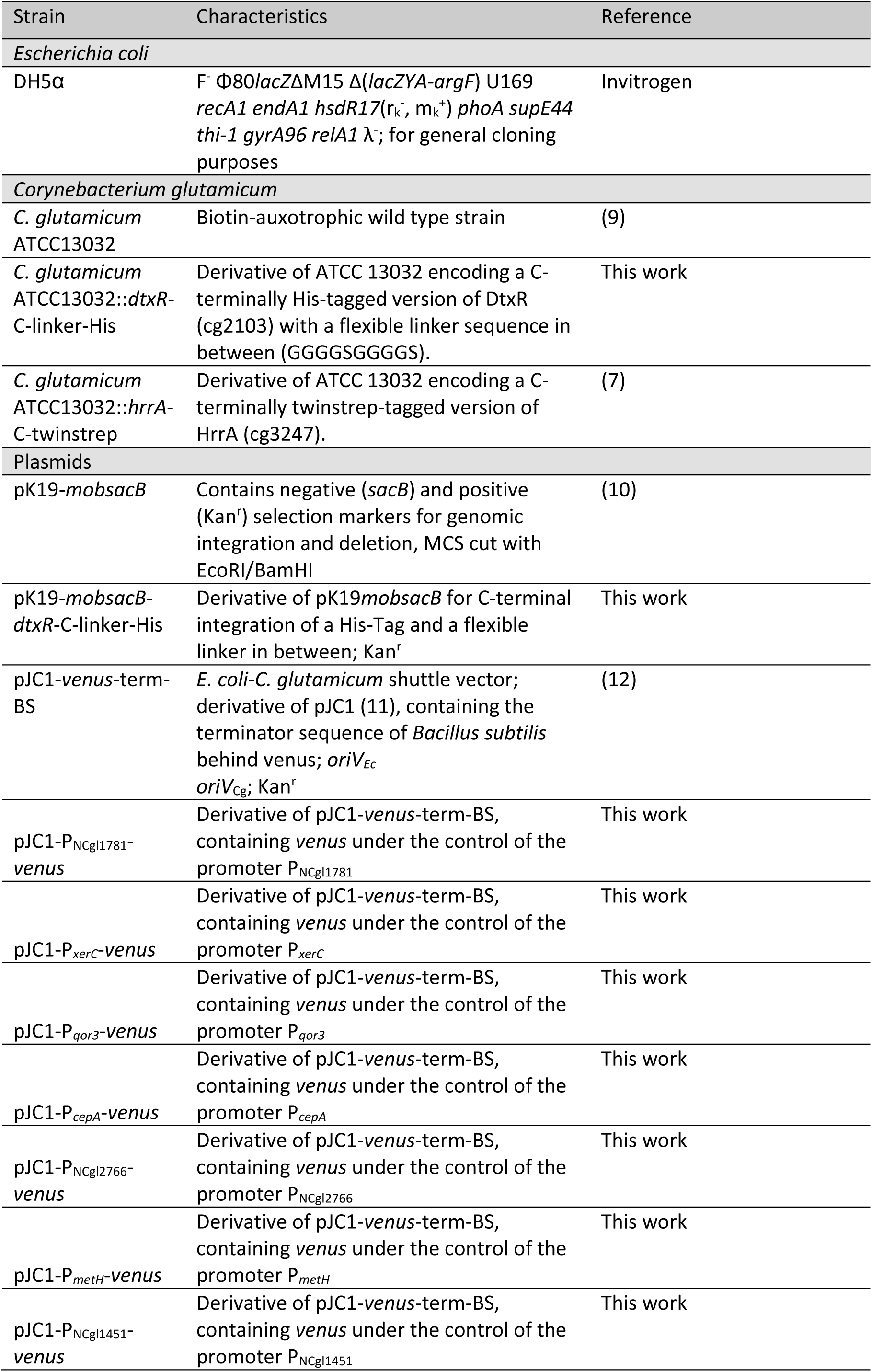
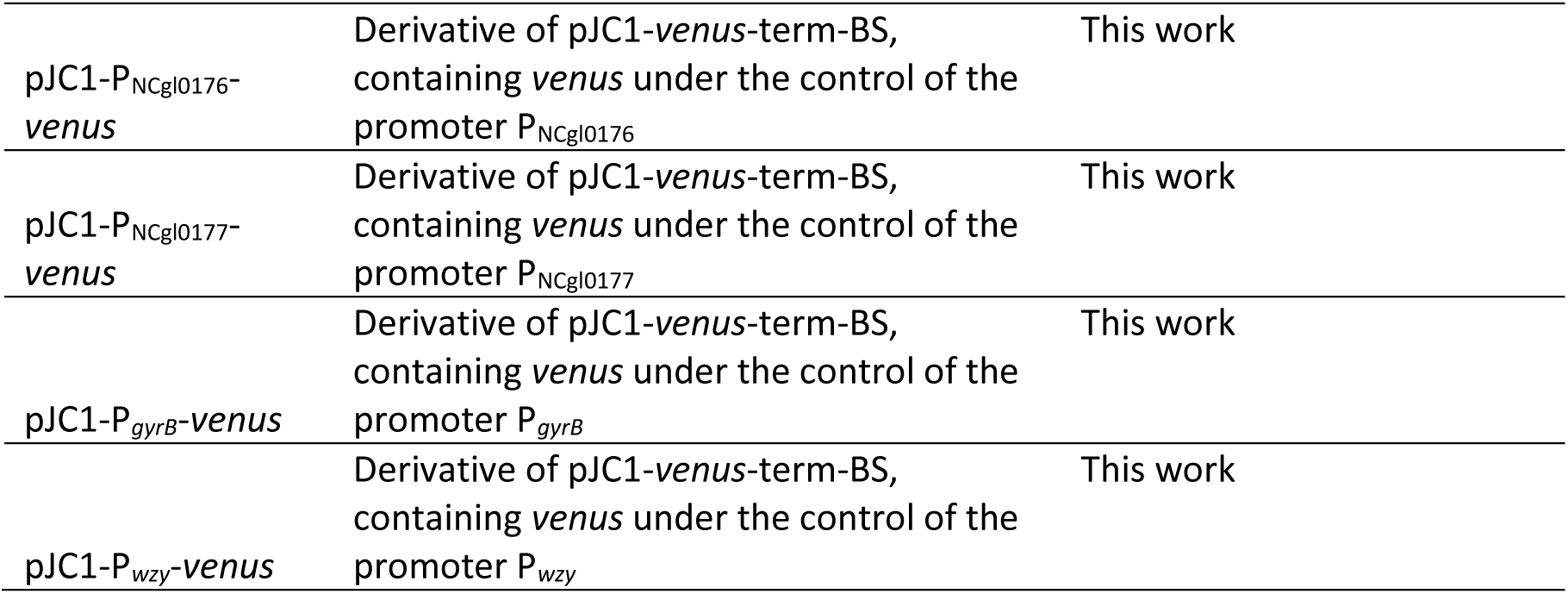
Bacterial strains and plasmids used in this study.

**Table S2:**
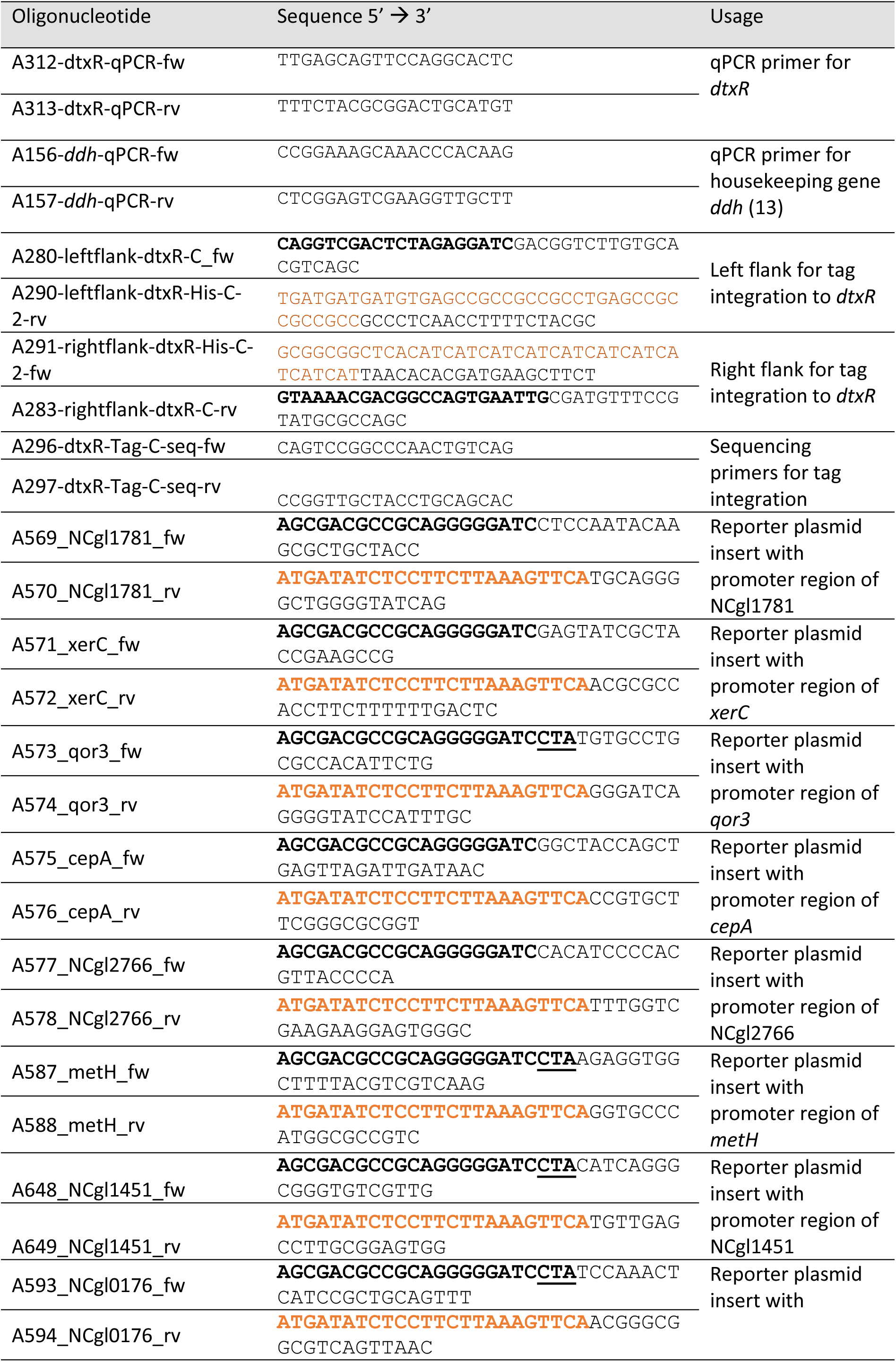
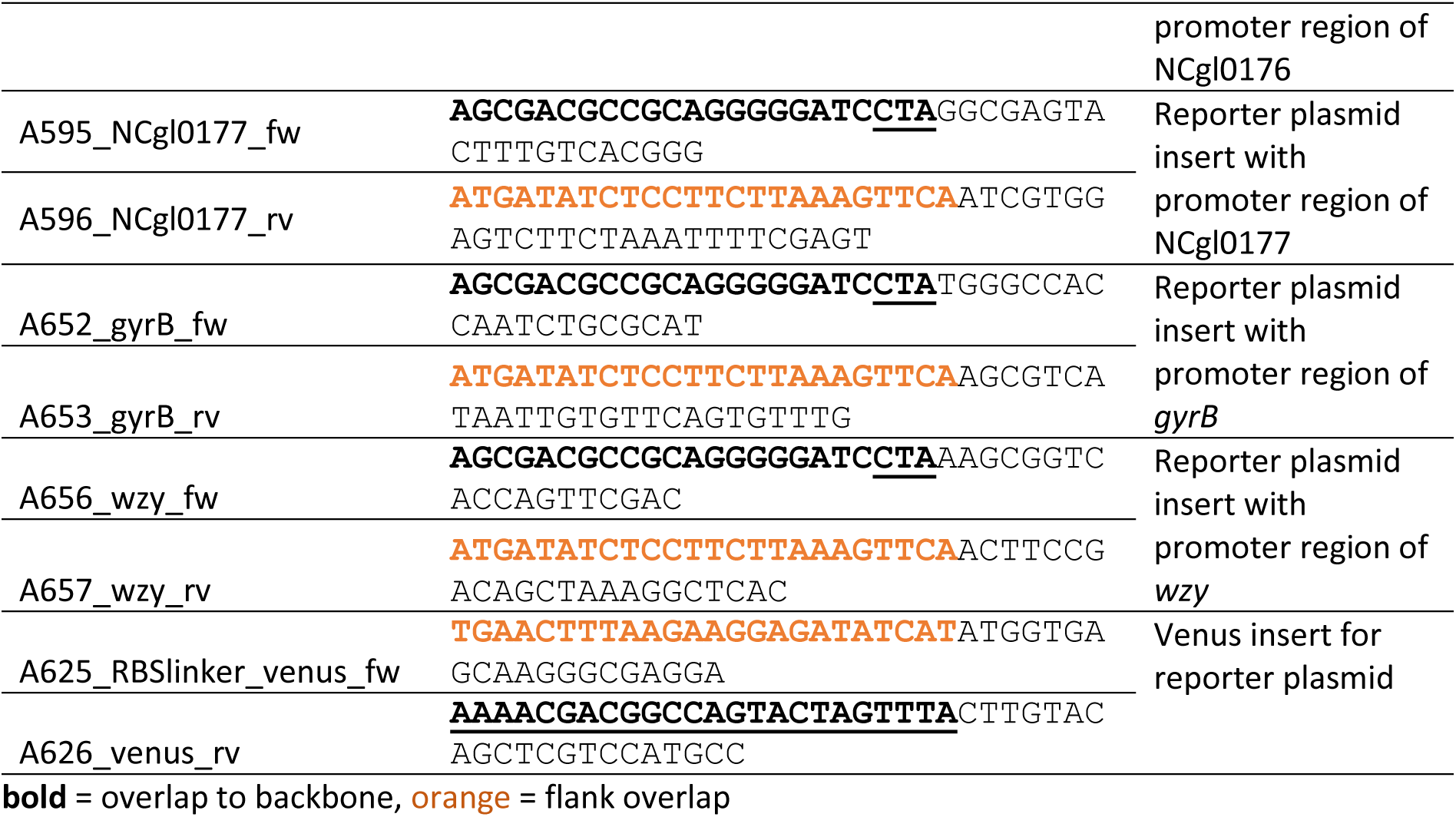
Oligonucleotides used in this study provided by Eurofins Genomics (Germany).

Table S3: Complete dataset of genome wide DtxR binding (ChAP-Seq) at iron excess (100 µM FeSO4) and heme (4 µM heme) conditions in triplicates. (external Excel file)

Table S4: Complete dataset of genome wide HrrA binding (ChAP-Seq) at iron excess (100 µM FeSO4) and heme (4 µM heme) conditions in triplicates. (external Excel file)

**Table S5:** Overlapping peaks for DtxR and HrrA.

| gene number | gene | DtxR iron excess |  | DtxR heme |  | HrrA iron excess |  | HrrA heme |  |
| --- | --- | --- | --- | --- | --- | --- | --- | --- | --- |
| | | $\mu$ peak intensity | n | $\mu$ peak intensity | n | $\mu$ peak intensity | n | $\mu$ peak intensity | n |
| NCgl0005 | *gyrB* | 2.01 | 1 | n.d. | 0 | 2.80 | 2 | n.d. | 0 |
| NCgl0177/<br><u>NCgl0176</u> |  | 2.03 | 1 | n.d. | 0 | 2.42 | 3 | 3.45 | 1 |
| NCgl0352 | *wzy* | 1.94 | 1 | n.d. | 0 | 5.44 | 3 | 7.71 | 3 |
| NCgl0353 |  | n.d. | 0 | 2.02 | 1 | n.d. | 0 | 3.38 | 1 |
| NCgl0358/<br>NCgl0359 | *ramB*/<br>*sdhCD* | 7.93 | 3 | 3.16 | 3 | 50.47 | 3 | 72.92 | 3 |
| NCgl1200 |  | 2.24 | 1 | n.d. | 0 | n.d. | 0 | 2.50 | 1 |
| <u>NCgl1395</u> |  | 3.31 | 3 | 2.56 | 3 | 3.07 | 3 | 3.75 | 3 |
| NCgl1444 |  | 2.23 | 2 | n.d. | 0 | 9.48 | 3 | 12.28 | 3 |
| NCgl1952 | *xerC* | n.d. | 0 | 6.54 | 3 | 43.06 | 3 | 254.09 | 3 |
| NCgl2146 | *hmuO* | 16.51 | 3 | 15.80 | 3 | 3.01 | 3 | 4.96 | 3 |
| <u>NCgl2766</u> |  | 2.29 | 3 | n.d. | 0 | 3.33 | 2 | n.d. | 0 |
| NCgl2834 | *hrrA* | n.d. | 0 | 3.19 | 2 | 2.99 | 3 | 6.29 | 3 |
| NCgl2842 | *uspA3* | n.d. | 0 | 2.02 | 1 | 3.64 | 2 | n.d. | 0 |
| NCgl2903/<br>NCgl2902 | *cepA*/<br>*qor3* | 2.53 | 2 | n.d. | 0 | n.d. | 0 | 3.11 | 3 |
| NCgl2968 |  | 1.89 | 1 | n.d. | 0 | 4.71 | 3 | 6.75 | 3 |
| NCgl2980 |  | n.d. | 0 | 2.00 | 1 | 2.25 | 2 | n.d. | 0 |

$\mu$ = mean value, n = number of replicates with a peak identified, n.d. = no peak detected, underlined = peak was annotated for HrrA for the gene into the other direction, i.e. binding probably not at same site.

## References

1. Andrews SC, Robinson AK, Rodríguez-Quiñones F. 2003. Bacterial iron homeostasis. FEMS Microbiol Rev 27:215–37 doi:10.1016/s0168-6445(03)00055-x.

2. Cornelis P, Wei Q, Andrews SC, Vinckx T. 2011. Iron homeostasis and management of oxidative stress response in bacteria. Metallomics 3:540–9 doi:10.1039/c1mt00022e.

3. Frunzke J, Gätgens C, Brocker M, Bott M. 2011. Control of heme homeostasis in *Corynebacterium glutamicum* by the two-component system HrrSA. Journal of Bacteriology 193:1212–21 doi:10.1128/jb.01130-10.

4. Heyer A, Gatgens C, Hentschel E, Kalinowski J, Bott M, Frunzke J. 2012. The two-component system ChrSA is crucial for haem tolerance and interferes with HrrSA in haem-dependent gene regulation in *Corynebacterium glutamicum*. Microbiology 158:3020–31 doi:10.1099/mic.0.062638-0.

5. Schmitt MP, Holmes RK. 1993. Analysis of diphtheria toxin repressor-operator interactions and characterization of a mutant repressor with decreased binding activity for divalent metals. Molecular Microbiology 9:173–181 10.1111/j.1365-2958.1993.tb01679.x.

6. Keppel M, Davoudi E, Gätgens C, Frunzke J. 2018. Membrane Topology and Heme Binding of the Histidine Kinases HrrS and ChrS in *Corynebacterium glutamicum*. Frontiers in Microbiology 9:183 doi:10.3389/fmicb.2018.00183.

7. Boyd J, Oza MN, Murphy JR. 1990. Molecular cloning and DNA sequence analysis of a diphtheria tox iron-dependent regulatory element (*dtxR*) from *Corynebacterium diphtheriae*. Proceedings of the National Academy of Sciences 87:5968–5972 doi:doi:10.1073/pnas.87.15.5968.

8. Pappenheimer AMJ, Johnson SJ. 1936. Studies in Diphtheria Toxin Production. I: The Effect of Iron and Copper. British Journal of Experimental Pathology 17:335–341.

9. Gold B, Rodriguez GM, Marras SA, Pentecost M, Smith I. 2001. The *Mycobacterium tuberculosis* IdeR is a dual functional regulator that controls transcription of genes involved in iron acquisition, iron storage and survival in macrophages. Mol Microbiol 42:851–65 doi:10.1046/j.1365-2958.2001.02684.x.

10. Ando M, Manabe YC, Converse PJ, Miyazaki E, Harrison R, Murphy JR, Bishai WR. 2003. Characterization of the role of the divalent metal ion-dependent transcriptional repressor MntR in the virulence of *Staphylococcus aureus*. Infect Immun 71:2584–90 doi:10.1128/iai.71.5.2584-2590.2003.

11. Wennerhold J, Bott M. 2006. The DtxR regulon of *Corynebacterium glutamicum*. Journal of Bacteriology 188:2907–18 doi:10.1128/jb.188.8.2907-2918.2006.

12. Brune I, Werner H, Hüser AT, Kalinowski J, Pühler A, Tauch A. 2006. The DtxR protein acting as dual transcriptional regulator directs a global regulatory network involved in iron metabolism of *Corynebacterium glutamicum*. BMC Genomics 7:21 doi:10.1186/1471-2164-7-21.

13. White A, Ding X, vanderSpek JC, Murphy JR, Ringe D. 1998. Structure of the metal-ion-activated diphtheria toxin repressor/ tox operator complex. Nature 394:502–506 doi:10.1038/28893.

14. Drazek ES, Hammack CA, Schmitt MP. 2000. *Corynebacterium diphtheriae* genes required for acquisition of iron from haemin and haemoglobin are homologous to ABC haemin transporters. Mol Microbiol 36:68–84 doi:10.1046/j.1365-2958.2000.01818.x.

15. Krüger A, Keppel M, Sharma V, Frunzke J. 2022. The diversity of heme sensor systems - heme- responsive transcriptional regulation mediated by transient heme protein interactions. FEMS Microbiol Rev doi:10.1093/femsre/fuac002 doi:10.1093/femsre/fuac002.

16. Hentschel E, Mack C, Gatgens C, Bott M, Brocker M, Frunzke J. 2014. Phosphatase activity of the histidine kinases ensures pathway specificity of the ChrSA and HrrSA two-component systems in *Corynebacterium glutamicum*. Molecular Microbiology 92:1326–42 doi:10.1111/mmi.12633.

17. Keppel M, Piepenbreier H, Gätgens C, Fritz G, Frunzke J. 2019. Toxic but tasty - temporal dynamics and network architecture of heme-responsive two-component signaling in *Corynebacterium glutamicum*. Mol Microbiol 111:1367–1381 doi:10.1111/mmi.14226.

18. Keppel M, Hünnefeld M, Filipchyk A, Viets U, Davoudi CF, Krüger A, Mack C, Pfeifer E, Polen T, Baumgart M, Bott M, Frunzke J. 2020. HrrSA orchestrates a systemic response to heme and determines prioritization of terminal cytochrome oxidase expression. Nucleic Acids Research 48:6547–6562 doi:10.1093/nar/gkaa415.

19. Kunkle CA, Schmitt MP. 2005. Analysis of a DtxR-regulated iron transport and siderophore biosynthesis gene cluster in *Corynebacterium diphtheriae*. J Bacteriol 187:422–33 doi:10.1128/jb.187.2.422-433.2005.

20. Kunkle CA, Schmitt MP. 2003. Analysis of the *Corynebacterium diphtheriae* DtxR regulon: identification of a putative siderophore synthesis and transport system that is similar to the *Yersinia* high-pathogenicity island-encoded yersiniabactin synthesis and uptake system. J Bacteriol 185:6826–40 doi:10.1128/jb.185.23.6826-6840.2003.

21. Keilhauer C, Eggeling L, Sahm H. 1993. Isoleucine synthesis in *Corynebacterium glutamicum*: molecular analysis of the *ilvB-ilvN-ilvC* operon. J Bacteriol 175:5595–603 doi:10.1128/jb.175.17.5595-5603.1993.

22. Kensy F, Zang E, Faulhammer C, Tan R-K, Büchs J. 2009. Validation of a high-throughput fermentation system based on online monitoring of biomass and fluorescence in continuously shaken microtiter plates. Microbial Cell Factories 8:31 doi:10.1186/1475-2859-8-31.

23. Sambrook JF, Russell D. 2001. Molecular Cloning: A Laboratory Manual (3-Volume Set), vol 1.

24. Eikmanns BJ, Thum-Schmitz N, Eggeling L, Lüdtke K-U, Sahm H. 1994. Nucleotide sequence, expression and transcriptional analysis of the *Corynebacterium glutamicum gltA* gene encoding citrate synthase. Microbiology 140:1817–1828 10.1099/13500872-140-8-1817.

25. Gibson DG, Young L, Chuang RY, Venter JC, Hutchison CA, 3rd, Smith HO. 2009. Enzymatic assembly of DNA molecules up to several hundred kilobases. Nat Methods 6:343–5 doi:10.1038/nmeth.1318.

26. Schäfer A, Tauch A, Jäger W, Kalinowski J, Thierbach G, Pühler A. 1994. Small mobilizable multi- purpose cloning vectors derived from the *Escherichia coli* plasmids pK18 and pK19: selection of defined deletions in the chromosome of *Corynebacterium glutamicum*. Gene 145:69–73 doi:10.1016/0378-1119(94)90324-7.

27. van der Rest ME, Lange C, Molenaar D. 1999. A heat shock following electroporation induces highly efficient transformation of *Corynebacterium glutamicum* with xenogeneic plasmid DNA. Applied Microbiology and Biotechnology 52:541–545 doi:10.1007/s002530051557.

28. Niebisch A, Bott M. 2001. Molecular analysis of the cytochrome bc1-aa3 branch of the *Corynebacterium glutamicum* respiratory chain containing an unusual diheme cytochrome c1. Arch Microbiol 175:282–94 doi:10.1007/s002030100262.

29. Frunzke J, Bramkamp M, Schweitzer JE, Bott M. 2008. Population Heterogeneity in *Corynebacterium glutamicum* ATCC 13032 caused by prophage CGP3. J Bacteriol 190:5111–9 doi:10.1128/jb.00310-08.

30. Livak KJ, Schmittgen TD. 2001. Analysis of relative gene expression data using real-time quantitative PCR and the 2^-ΔΔCt^ Method. Methods 25:402–8 doi:10.1006/meth.2001.1262.

31. Pfeifer E, Hünnefeld M, Popa O, Polen T, Kohlheyer D, Baumgart M, Frunzke J. 2016. Silencing of cryptic prophages in *Corynebacterium glutamicum*. Nucleic acids research 44:10117–10131 doi:10.1093/nar/gkw692.

32. Langmead B, Salzberg SL. 2012. Fast gapped-read alignment with Bowtie 2. Nat Methods 9:357–9 doi:10.1038/nmeth.1923.

33. Langmead B, Wilks C, Antonescu V, Charles R. 2019. Scaling read aligners to hundreds of threads on general-purpose processors. Bioinformatics 35:421–432 doi:10.1093/bioinformatics/bty648.

34. Chen X, Zaro JL, Shen WC. 2013. Fusion protein linkers: property, design and functionality. Adv Drug Deliv Rev 65:1357–69 doi:10.1016/j.addr.2012.09.039.

35. Layer G. 2021. Heme biosynthesis in prokaryotes. Biochim Biophys Acta Mol Cell Res 1868:118861 doi:10.1016/j.bbamcr.2020.118861.

36. Wilks A, Schmitt MP. 1998. Expression and Characterization of a Heme Oxygenase (HmuO) from *Corynebacterium diphtheriae*: Iron Acquisition Requires Oxidative Cleavage of the Heme Macrocycle. Journal of Biological Chemistry 273:837–841 doi:10.1074/jbc.273.2.837.

37. Bailey TL, Elkan C. 1994. Fitting a mixture model by expectation maximization to discover motifs in biopolymers. Proc Int Conf Intell Syst Mol Biol 2:28–36.

38. Madeira F, Madhusoodanan N, Lee J, Eusebi A, Niewielska A, Tivey ARN, Lopez R, Butcher S. 2024. The EMBL-EBI Job Dispatcher sequence analysis tools framework in 2024. Nucleic acids research 52:W521–W525 doi:10.1093/nar/gkae241.

39. Grant CE, Bailey TL, Noble WS. 2011. FIMO: scanning for occurrences of a given motif. Bioinformatics 27:1017–1018 doi:10.1093/bioinformatics/btr064.

40. Escorcia-Rodríguez J, Tauch A, Freyre-González J. 2020. Corynebacterium glutamicum Regulation beyond Transcription: Organizing Principles and Reconstruction of an Extended Regulatory Network Incorporating Regulations Mediated by Small RNA and Protein-Protein Interactions doi:10.20944/preprints202012.0503.v1. Preprints.org.

41. Pi H, Helmann JD. 2017. Sequential induction of Fur-regulated genes in response to iron limitation in *Bacillus subtilis*. Proc Natl Acad Sci U S A 114:12785–12790 doi:10.1073/pnas.1713008114.

42. Pi H, Helmann JD. 2018. Genome-Wide Characterization of the Fur Regulatory Network Reveals a Link between Catechol Degradation and Bacillibactin Metabolism in Bacillus subtilis. mBio 9 doi:10.1128/mBio.01451-18.

43. Myers KS, Yan H, Ong IM, Chung D, Liang K, Tran F, Keleş S, Landick R, Kiley PJ. 2013. Genome- scale analysis of *Escherichia coli* FNR reveals complex features of transcription factor binding. PLoS Genet 9:e1003565 doi:10.1371/journal.pgen.1003565.

44. Myers KS, Park DM, Beauchene NA, Kiley PJ. 2015. Defining bacterial regulons using ChIP-seq. Methods 86:80–8 doi:10.1016/j.ymeth.2015.05.022.

45. Rückert C, Pühler A, Kalinowski J. 2003. Genome-wide analysis of the l-methionine biosynthetic pathway in *Corynebacterium glutamicum* by targeted gene deletion and homologous complementation. Journal of Biotechnology 104:213–228 10.1016/S0168-1656(03)00158-5.

46. Yellaboina S, Ranjan S, Chakhaiyar P, Hasnain SE, Ranjan A. 2004. Prediction of DtxR regulon: identification of binding sites and operons controlled by Diphtheria toxin repressor in Corynebacterium diphtheriae. BMC Microbiol 4:38 doi:10.1186/1471-2180-4-38.

47. Stojiljkovic I, Bäumler AJ, Hantke K. 1994. Fur Regulon in Gram-negative Bacteria: Identification and Characterization of New Iron-regulated *Escherichia coli* Genes by a Fur Titration Assay. Journal of Molecular Biology 236:531–545 10.1006/jmbi.1994.1163.

48. Levine RL, Mosoni L, Berlett BS, Stadtman ER. 1996. Methionine residues as endogenous antioxidants in proteins. Proc Natl Acad Sci U S A 93:15036–40 doi:10.1073/pnas.93.26.15036.

49. Slyshenkov VS, Shevalye AA, Liopo AV, Wojtczak L. 2002. Protective role of L-methionine against free radical damage of rat brain synaptosomes. Acta Biochim Pol 49:907–16.

50. Brot N, Weissbach H. 1983. Biochemistry and physiological role of methionine sulfoxide residues in proteins. Arch Biochem Biophys 223:271–81 doi:10.1016/0003-9861(83)90592-1.

51. Si M, Zhang L, Chaudhry MT, Ding W, Xu Y, Chen C, Akbar A, Shen X, Liu SJ. 2015. *Corynebacterium glutamicum* methionine sulfoxide reductase A uses both mycoredoxin and thioredoxin for regeneration and oxidative stress resistance. Appl Environ Microbiol 81:2781–96 doi:10.1128/aem.04221-14.

52. Erdmann K, Grosser N, Schroder H. 2005. L-methionine reduces oxidant stress in endothelial cells: Role of heme oxygenase-1, ferritin, and nitric oxide. The AAPS journal 7:E195–200 doi:10.1208/aapsj070118.

53. Grindley ND, Whiteson KL, Rice PA. 2006. Mechanisms of site-specific recombination. Annu Rev Biochem 75:567–605 doi:10.1146/annurev.biochem.73.011303.073908.

54. Blakely G, Colloms S, May G, Burke M, Sherratt D. 1991. *Escherichia coli* XerC recombinase is required for chromosomal segregation at cell division. New Biol 3:789–98.

55. Aussel L, Barre F-X, Aroyo M, Stasiak A, Stasiak AZ, Sherratt D. 2002. FtsK Is a DNA Motor Protein that Activates Chromosome Dimer Resolution by Switching the Catalytic State of the XerC and XerD Recombinases. Cell 108:195–205 10.1016/S0092-8674(02)00624-4.

56. Seo SW, Kim D, Latif H, O’Brien EJ, Szubin R, Palsson BO. 2014. Deciphering Fur transcriptional regulatory network highlights its complex role beyond iron metabolism in *Escherichia coli*. Nature Communications 5:4910 doi:10.1038/ncomms5910.

57. Helfrich S, Pfeifer E, Krämer C, Sachs CC, Wiechert W, Kohlheyer D, Nöh K, Frunzke J. 2015. Live cell imaging of SOS and prophage dynamics in isogenic bacterial populations. Molecular Microbiology 98:636–650 10.1111/mmi.13147.

58. Boeselager R, Pfeifer E, Frunzke J. 2018. Cytometry meets next-generation sequencing – RNA-Seq of sorted subpopulations reveals regional replication and iron-triggered prophage induction in *Corynebacterium glutamicum*. Scientific Reports 8 doi:10.1038/s41598-018-32997-9.

59. Dos Santos NM, Picinato BA, Santos LS, de Araújo HL, Balan A, Koide T, Marques MV. 2024. Mapping the IscR regulon sheds light on the regulation of iron homeostasis in *Caulobacter*. Front Microbiol 15:1463854 doi:10.3389/fmicb.2024.1463854.

60. Galagan JE, Minch K, Peterson M, Lyubetskaya A, Azizi E, Sweet L, Gomes A, Rustad T, Dolganov G, Glotova I, Abeel T, Mahwinney C, Kennedy AD, Allard R, Brabant W, Krueger A, Jaini S, Honda B, Yu WH, Hickey MJ, Zucker J, Garay C, Weiner B, Sisk P, Stolte C, Winkler JK, Van de Peer Y, Iazzetti P, Camacho D, Dreyfuss J, Liu Y, Dorhoi A, Mollenkopf HJ, Drogaris P, Lamontagne J, Zhou Y, Piquenot J, Park ST, Raman S, Kaufmann SH, Mohney RP, Chelsky D, Moody DB, Sherman DR, Schoolnik GK. 2013. The *Mycobacterium tuberculosis* regulatory network and hypoxia. Nature 499:178–83 doi:10.1038/nature12337.

61. MacPhillamy C, Alinejad-Rokny H, Pitchford WS, Low WY. 2022. Cross-species enhancer prediction using machine learning. Genomics doi:10.1016/j.ygeno.2022.110454:110454 doi:10.1016/j.ygeno.2022.110454.

62. Qin S, Yuan Y, Huang X, Tan Z, Hu X, Liu H, Pu Y, Ding YQ, Su Z, He C. 2022. Topoisomerase IIA in adult NSCs regulates SVZ neurogenesis by transcriptional activation of Usp37. Nucleic Acids Res doi:10.1093/nar/gkac731 doi:10.1093/nar/gkac731.

63. Rustad TR, Minch KJ, Ma S, Winkler JK, Hobbs S, Hickey M, Brabant W, Turkarslan S, Price ND, Baliga NS, Sherman DR. 2014. Mapping and manipulating the *Mycobacterium tuberculosis* transcriptome using a transcription factor overexpression-derived regulatory network. Genome Biology 15:502 doi:10.1186/s13059-014-0502-3.

64. Sims PA, Wong CF, Vuga D, McCammon JA, Sefton BM. 2005. Relative contributions of desolvation, inter- and intramolecular interactions to binding affinity in protein kinase systems. Journal of Computational Chemistry 26:668–681 10.1002/jcc.20207.

65. Ponta H, Cato AC, Herrlich P. 1992. Interference of pathway specific transcription factors. Biochim Biophys Acta 1129:255–61 doi:10.1016/0167-4781(92)90501-p.

66. Dorman CJ. 2019. DNA supercoiling and transcription in bacteria: a two-way street. BMC Molecular and Cell Biology 20:26 doi:10.1186/s12860-019-0211-6.

## References

1. Kensy F, Zang E, Faulhammer C, Tan R-K, Büchs J. 2009. Validation of a high-throughput fermentation system based on online monitoring of biomass and fluorescence in continuously shaken microtiter plates. Microb Cell Factories 8:31 doi:10.1186/1475-2859-8-31.

2. Livak KJ, Schmittgen TD. 2001. Analysis of relative gene expression data using real-time quantitative PCR and the 2^-ΔΔCt^ Method. Methods 25:402–8 doi:10.1006/meth.2001.1262.

3. Wennerhold J, Bott M. 2006. The DtxR regulon of *Corynebacterium glutamicum*. Journal of Bacteriology 188:2907–18 doi:10.1128/jb.188.8.2907-2918.2006.

4. Brune I, Werner H, Hüser AT, Kalinowski J, Pühler A, Tauch A. 2006. The DtxR protein acting as dual transcriptional regulator directs a global regulatory network involved in iron metabolism of *Corynebacterium glutamicum*. BMC Genomics 7:21 doi:10.1186/1471-2164-7-21.

5. Madeira F, Madhusoodanan N, Lee J, Eusebi A, Niewielska A, Tivey ARN, Lopez R, Butcher S. 2024. The EMBL-EBI Job Dispatcher sequence analysis tools framework in 2024. Nucleic acids research 52:W521–W525 doi:10.1093/nar/gkae241.

6. Bailey TL, Elkan C. 1994. Fitting a mixture model by expectation maximization to discover motifs in biopolymers. Proc Int Conf Intell Syst Mol Biol 2:28–36.

7. Keppel M, Hünnefeld M, Filipchyk A, Viets U, Davoudi CF, Krüger A, Mack C, Pfeifer E, Polen T, Baumgart M, Bott M, Frunzke J. 2020. HrrSA orchestrates a systemic response to heme and determines prioritization of terminal cytochrome oxidase expression. Nucleic Acids Research 48:6547–6562 doi:10.1093/nar/gkaa415.

8. Grant CE, Bailey TL, Noble WS. 2011. FIMO: scanning for occurrences of a given motif. Bioinformatics 27:1017–1018 doi:10.1093/bioinformatics/btr064.

9. Kinoshita S, Udaka S, Shimono M. 2004. Studies on the amino acid fermentation. Part 1. Production of L-glutamic acid by various microorganisms. The Journal of General and Applied Microbiology 50:331–43.

10. Schäfer A, Tauch A, Jäger W, Kalinowski J, Thierbach G, Pühler A. 1994. Small mobilizable multi-purpose cloning vectors derived from the *Escherichia coli* plasmids pK18 and pK19: selection of defined deletions in the chromosome of *Corynebacterium glutamicum*. Gene 145:69–73 doi:10.1016/0378-1119(94)90324-7.

11. Cremer J, Eggeling L, Sahm H. 1990. Cloning the *dapA dapB* cluster of the lysine-secreting bacterium *Corynebacterium glutamicum*. Molecular and General Genetics MGG 220:478–480 doi:10.1007/BF00391757.

12. Baumgart M, Luder K, Grover S, Gätgens C, Besra GS, Frunzke J. 2013. IpsA, a novel LacI-type regulator, is required for inositol-derived lipid formation in *Corynebacteria* and *Mycobacteria*. BMC Biol 11:122 doi:10.1186/1741-7007-11-122.

13. Frunzke J, Bramkamp M, Schweitzer JE, Bott M. 2008. Population Heterogeneity in *Corynebacterium glutamicum* ATCC 13032 caused by prophage CGP3. J Bacteriol 190:5111–9 doi:10.1128/jb.00310-08.

